# Mitochondrial genes in the 22q11.2 deleted region regulate neural stem and progenitor cell proliferation

**DOI:** 10.1101/2023.01.03.522615

**Authors:** Philip D. Campbell, Isaiah Lee, Summer Thyme, Michael Granato

**Author notes:** Corresponding Author: Michael Granato.

## Abstract

Microdeletion of a 3Mbp region encompassing 45 protein-coding genes at chromosome 22q11.2 (22q11.2DS) predisposes to multiple neurodevelopmental disorders and is one of the greatest genetic risk factors for schizophrenia. Defective mitochondrial function has been hypothesized to contribute to 22q11.2DS pathogenesis; however, which of the six mitochondrial genes contribute to neurodevelopmental phenotypes and their underlying mechanisms remain unresolved. To systematically test 22q11.2DS genes for functional roles in neurodevelopment and behavior, we generated genetic mutants for each of the 37 conserved zebrafish orthologs and performed high throughput behavioral phenotyping using seven behavioral assays. Through this unbiased approach, we identified five single-gene mutants with partially overlapping behavioral phenotypes. Two of these genes, *mrpl40* and *prodha*, encode for mitochondrial proteins and, similar to what we observed in *mrpl40* and *prodha* mutants, pharmacologic inhibition of mitochondrial function during development results in microcephaly. Finally, we show that both *mrpl40* and *prodha* mutants display neural stem and progenitor cell phenotypes, with each gene regulating different neural stem cell populations. Combined, our results demonstrate a critical role for mitochondrial function in neural stem and progenitor cell populations in the developing vertebrate brain and provide compelling evidence that mitochondrial dysfunction during neurodevelopment is linked to brain volume and behavioral phenotypes observed in models of 22q11.2DS.

## Introduction

Heterozygous microdeletion at chromosome 22q11.2 (22q11.2DS) is the most common microdeletion syndrome, occurring in 1 out of every ∼2000 live births^1^. 22q11.2DS predisposes individuals to multiple neurodevelopmental disorders (NDDs), including schizophrenia (SZ), autism spectrum disorders (ASD), intellectual disability (ID), and attention-deficit/hyperactivity disorder (ADHD)^2^. Mouse models with similar heterozygous deletions display developmental deficits in neurogenesis^3^, brain size^4^, and ultimately circuit-level^5^ as well as behavioral abnormalities^6^, suggesting that this genetic locus has an integral role in early brain development that impacts postnatal behaviors and intellectual abilities. Thus, understanding the molecular mechanisms by which 22q11.2DS genes regulate brain development and behavior is fundamental to identifying therapeutic targets for individuals with 22q11.2DS and idiopathic forms of NDDs. Yet despite their importance, the neurodevelopmental and behavioral roles of all genes within the deleted region remain incompletely understood.

In the majority of cases, the deleted region encompasses 45 protein coding genes, of which ∼90% are expressed in the brain in humans^7^, suggesting that many genes within the region have the potential to contribute to neurodevelopmental phenotypes. While a subset of genes have been implicated in neuronal function and behavior, data on brain and/or behavioral phenotypes of individual mouse knockouts exists for only a fraction of the 45 genes^8^. Even for those available, homozygous knockouts are frequently embryonic lethal, which further complicates the analysis of functional roles these genes might have during later developmental timepoints. Finally, a prevailing yet not fully tested hypothesis is that rather than single genes driving disease, interactions between genes in the region that overlap in function could be a driving factor^9^, adding to the complexity of disentangling the functional contributions of the 45 genes to neurodevelopmental and behavioral phenotypes.

Recent work has suggested that genes with mitochondrial function may be one such driving factor^10–15^. Indeed, proteins encoded by at least six genes in the deleted region are localized to mitochondria^10^ and neurons derived from individuals with the deletion display deficits in mitochondrial function^11^. Patient phenotypic severity also appears to be correlated with deficits in mitochondrial function in patient cells lines^12^. At the same time, multiple lines of evidence over the last decade have implicated mitochondria in neurogenesis^16^. In fact, neurogenesis is affected in deletion mice^3^ and individuals with 22q11.2DS display microcephaly, yet whether 22q11.2DS mitochondrial genes contribute to these phenotypes is an unresolved question.

To systematically test 22q11.2DS genes for their roles in neurodevelopment and behavior, we generated presumptive null mutants for all 37 conserved 22q11.2DS zebrafish orthologs. We then subjected these mutants to seven validated high-throughput larval behavioral assays that measure motor behavior, sensorimotor processing, habituation learning, and sensorimotor gating. This revealed five genes with partially overlapping behavioral phenotypes, suggesting roles in similar biologic processes and/or circuits. Two of these genes, *mrpl40* and *prodha*, encode mitochondrial genes, and similar to what we observe in these two mutants, pharmacologic inhibition of mitochondrial function leads to microcephaly. Moreover, we find that both *mrpl40* and *prodha* mutants display phenotypes in neural stem and progenitor cell (NSPC) populations. While both *mrpl40* and *prodha* mutants display abnormal NSPC proliferation, each gene disrupts distinct NSPC populations; *mrpl40* mutants display prominent phenotypes in highly proliferative progenitors, whereas *prodha* mutants display phenotypes in less proliferative radial glia-like cells. Finally, we show that *mrpl40*/*prodha* double mutants display more severe behavioral phenotypes than single mutants. Combined, these results identify two mitochondrial genes within the 22q11.2DS deleted region that play distinct roles in regulating brain size and NSPC proliferation *in vivo*. Furthermore, our results suggest that in addition to deleterious effects in post-mitotic neurons, defective mitochondrial functioning plays a role during neurogenesis in 22q11.2DS-linked pathogenesis earlier than previously thought. Together, these results reveal a clear functional link between mitochondrial gene dysfunction, microcephaly, and 22q11.2DS-relevant behavioral phenotypes.

## Results

### Generation of zebrafish 22q11.2DS orthologous gene mutants

To identify the protein-coding genes in the 22q11.2DS deleted region that play critical roles in neurodevelopment and behavior *in vivo*, we used the zebrafish model system. We previously reported that 37 of the 45 protein-coding genes in the 22q11.2DS deleted region have orthologs in zebrafish^17^, with seven having duplicate copies and one having triplicate copies for a total of 46 zebrafish 22q11.2DS orthologous genes (Figures 1A-1B). To generate presumptive null alleles for each of the individual 46 genes, we used a CRISPR-Cas9 targeting approach with two guide RNAs (gRNA) targeting regions ∼50-400bp apart, and flanking either a known functional domain or a highly conserved region (Table S1). Using this approach, we identified heterozygous F2 adults with a variety of insertions and deletions (Table S1). Mutations were predicted to lead to frameshifts (30/46, 65%), in-frame insertions/deletions (11/46, 24%), or disruption of splice sites (5/46, 11%) (Table S1). Five of the 46 mutant lines displayed gross morphologic defects in the homozygous state at 6 days post-fertilization (6dpf) (Figure 1B, Table S2), and these homozygous mutants were excluded from subsequent behavioral analyses. To assess behavior, we used a behavioral analysis pipeline that consisted of seven previously validated assays (described in detail in the following section). This pipeline identified five mutant lines that displayed behavioral defects at 6dpf (Figures 1B, 2). Importantly, these mutants did not display overt morphological defects, though two (*dgcr8* and *snap29*) had deficits in swim bladder inflation that were incompletely penetrant (Table S3). Thus, our single gene mutational analysis identified 10 genes with roles in either development or behavior prior to 6dpf.

**Figure 1.**
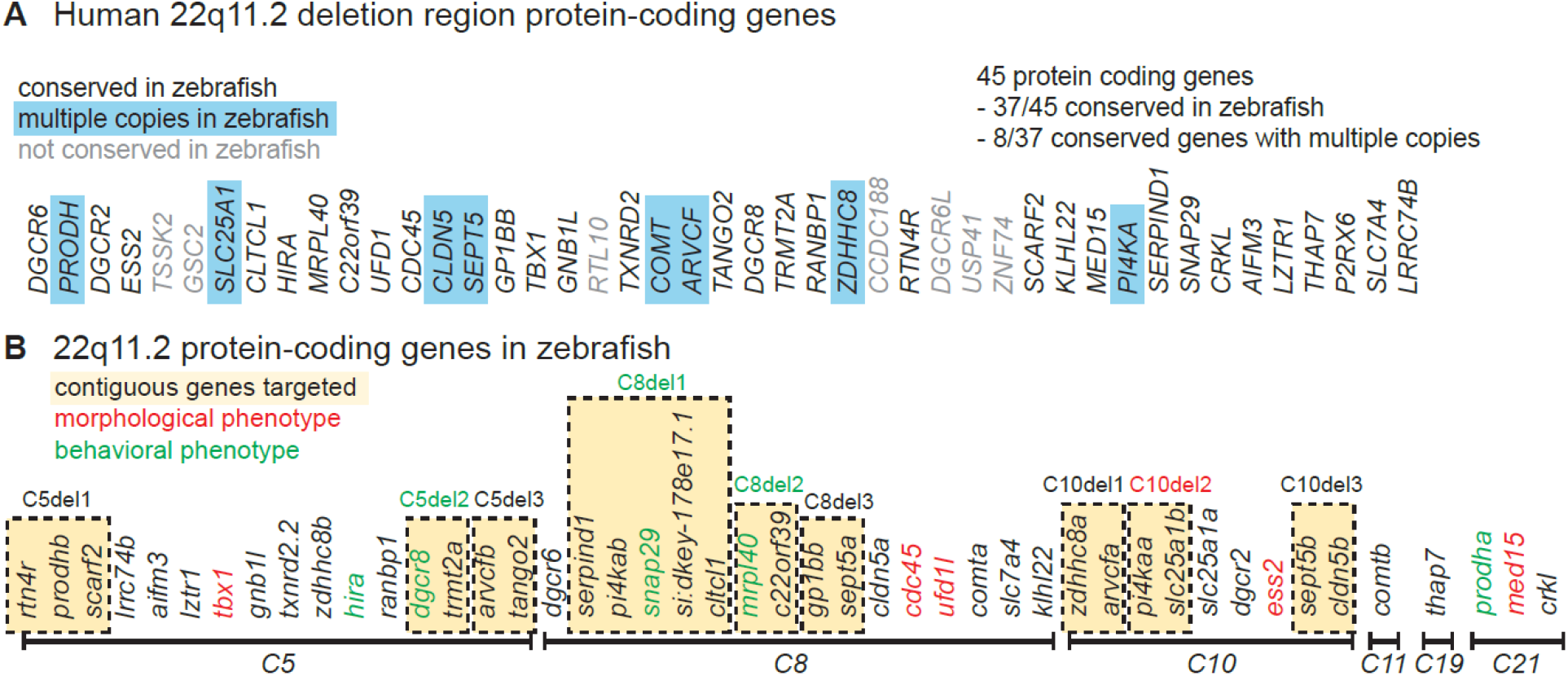
Generation of zebrafish 22q11.2DS orthologous gene mutants. (A) Protein-coding genes present in the usual 3Mbp human deletion. Those without orthologs in zebrafish are shaded gray. Those with multiple copies in zebrafish are highlighted in blue. (B)Zebrafish 22q11.2 orthologs are on six different chromosomes. Contiguous genes that were targeted for combination deletion are highlighted in yellow. Single or combination gene mutants with morphological or behavioral phenotypes are shown in red and green font respectively.

### Mutants in five genes including two encoding mitochondrial proteins, display partially overlapping behavioral phenotypes

As a first approach to identify genes with roles in brain development and/or function, we subjected all 46 single-gene mutant lines to behavioral phenotyping at 6dpf (Figure 2A). We used an unbiased phenotyping approach, assaying behavioral responses to visual and acoustic stimuli. Specifically, using visual stimuli, we assayed the visual motor response (VMR)^18^, the response to flashes of light (Light Flash, LF)^19^, the response to flashes of darkness (Dark Flash, DF)^19^, and Short-term habituation (Hab) of the DF response^20^. Using acoustic stimuli, we assayed the acoustic startle response (ASR)^21^, as well as ASR modulation, including startle sensitivity^22^, pre-pulse inhibition (PPI)^23^ and startle habituation^20^. Behaviors of individual larvae were tracked at high temporal resolution, and all larvae were genotyped following completion of the behavioral assay (Figure 2A). From each behavioral assay multiple kinematic parameters were computed using custom-written tracking software: frequency, features of movement (speed, distance traveled, etc.), and position metrics were computed for the VMR assay and frequency of response to stimulus and features of the responses (bend angle, latency, distance, etc.) were computed for LF, DF and ASR. Homozygous mutant larvae were compared to sibling larvae across all behavioral metrics. In cases where gross morphological defects were observed in homozygotes, heterozygotes were compared to wildtypes.

**Figure 2.**
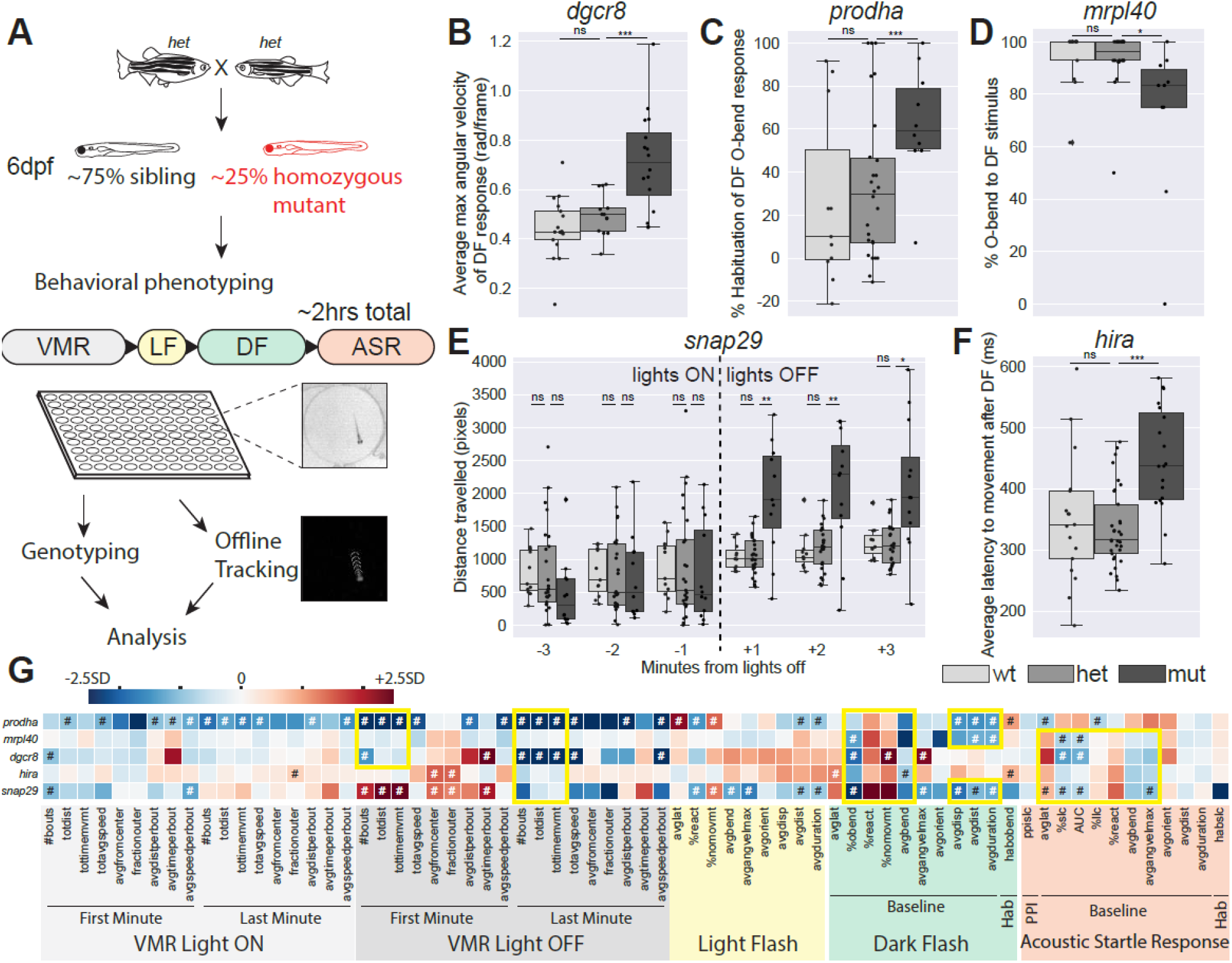
Mutants in five genes including two encoding mitochondrial proteins, display partially overlapping behavioral phenotypes. (A) Behavioral phenotyping paradigm. Heterozygous carriers are in-crossed to generate homozygous mutants and siblings that are raised to 6dpf at which point behavioral phenotyping is performed blinded to genotype. Approximately 50 larvae are used for each mutant line, and arrayed in a 100-well plate. Assay occurs over approximately 2 hours in the following order: Visual-motor Response (VMR), Light Flash (LF), Dark Flash (DF), Acoustic Startle Response (ASR). Individual larvae are genotyped, acquired videos are tracked offline, and then analyzed for 94 behavioral metrics. (B-F) Example phenotypes for five single gene mutants with reproducible behavioral phenotypes. Each plot represents data from a single behavioral run. Points indicate individual larvae, Boxes indicate range of 25^th^ to 75^th^ percentile with line indicating the median, whiskers indicate the rest of the range, with outliers lying outside of this. (B) Average maximum angular velocity of the DF response is increased in *dgcr8* mutants compared to siblings. (C) Percent habituation of DF O-bend response is increased in *prodha* mutants compared to siblings. (D) Percent of time *mrpl40* mutants perform O-bends in response to DF is decreased compared to siblings. (E) *snap29* mutants display an increase in distance travelled in VMR assay following lights turning off compared to siblings. Each bin represents the total distance travelled in during that minute of the assay. (F) Average latency to movement after DF stimulus is increased in *hira* mutants compared to siblings. All statistics represent one-way ANOVA, *p<0.05, **p<0.01, ***p<0.001. (G) Heatmap illustrating single gene mutant phenotypes across behavioral metrics. Each row is an individual mutant line and all columns represent a behavioral metric. Color of each box represents difference of average mutant value compared to siblings expressed as number of standard deviations (SD). Blue boxes indicate the value is lower in the mutant and Red boxes indicate the value is higher in the mutant. Mutant values that are significantly different from siblings based on Student’s t-test with a Bonferroni-corrected p-value of <0.05/94 are designated with #. Yellow boxes indicate behavioral similarities across mutant lines. VMR behavioral metrics are highlighted in gray, LF in yellow, DF in green, and ASR in orange. Supplemental Tables 5-7 diplay behavioral data for mutant lines with reproducible behavioral phenotypes. Python tracking scripts are in Supplemental Zip file 1.

Using this pipeline, we identified five single-gene mutants with reproducible behavioral phenotypes. Each mutant line had behavioral phenotypes that spanned more than one assay. In each case, phenotypes were undectable in heterozygotes which appeared indistinguishable from wild-type siblings. Examples of phenotypes are displayed in Figures 2B-2F. Closer examination and comparison of behavioral heatmaps revealed several overlapping phenotypes across mutant lines (Figure 2G). All mutant lines displayed abnormal baseline metrics for the DF response, indicating that these genes have roles in mediating sensorimotor behaviors in response to darkness. Three of the five lines (*mrpl40, dgcr8, snap29*) also displayed reductions in the sensitivity of the ASR response, while the fifth line (*prodha*) displayed differences in the opposite direction indicating these genes play roles in the hindbrain startle circuit or in downstream effectors. Consistent with phenotypic overlap, the five genes also share similarities in terms of their reported biological function: *prodha* and *mrpl40* have roles in mitochondria, while *dgrc8* and *hira* have roles in gene regulation. In addition to shared phenotypes, we also observed phenotypes unique to one or two mutant lines. For example, *snap29* mutants had increased movement metrics during the first minute of the VMR following lights turning off (Figure 2E), while *prodha* (Figure 2C) and *hira* mutants had increased habituation of the DF response, and *dgcr8* mutants displayed increased angular velocity of the DF response (Figure 2F). In summary, our systematic analysis of the 22q11.2DS deleted region identified five single genes that function to either establish or maintain several overlapping behaviors, consistent with the idea that these genes play roles in similar biological or circuit-level processes.

One hypothesis for how the 22q11.2DS deleted region generates disease risk is that genes within the region are involved in overlapping biological processes^9^ resulting in partial functional redundancy, such that inactivating individual genes fails to reveal measurable phenotypes. To assess whether the combined loss of multiple 22q11.2DS genes results in behavioral phenotypes, we used multiple gRNAs to generate nine combination gene mutants that disrupted between 2 and 5 neighboring genes (Figure 1B). This yielded deletions ranging in size from ∼7kbp-544kbp (Table S4). One of these deletions, *C10del2*, encompassed two genes, *pi4kaa* and *slc25a1b*, and resulted at 6dpf in gross morphologic defects not present in single deletion mutants of either of the two genes (Figure 1B, Table S2), indicating that *pi4kaa* and *slc25a1b* may have partially overlapping function during early development. We then analyzed behavior of combination mutant lines. While three of the nine combination mutant lines had reproducible behavioral phenotypes (*C5del2, C8del1, C8del2*), the phenotypes were also detectable in single-gene mutants in genes present in the deleted region of the combination lines (*dgcr8, snap29, mrpl40* respectively) (Figure 1B). Comparing behavioral heatmaps of these combination mutants with their corresponding single mutants revealed similar phenotypes (Figure S1), indicating that the loss of a single gene is driving phenotypes in these combination mutant lines. Although we only tested a limited number of multiple gene deletion combinations, our results indicate that syntenic 22qDS zebrafish orthologs have limited functional redundancy.

### Pharmacologic inhibition of mitochondrial function phenocopies the *mrpl40* mutant phenotype

Two of the five genes identified through our behavioral screening (*mrpl40* and *prodha*) encode for proteins that localize to mitochondria^10^, and defective mitochondrial function has been previously reported in the context of 22q11.2DS^11^ as well as in neurodevelopmental disorders associated with 22q11.2DS^24^. As such, we hypothesized that disrupted mitochondrial functioning might contribute to the behavioral phenotypes we observed in *mrpl40* and *prodha* mutants. *mrpl40* is a component of the mitochondrial ribosome, and thus its disruption would be expected to decrease mitochondrial translation and subsequently reduce protein levels of electron transport chain components. To test whether inhibiting mitochondrial translation affects behavior and recapitulates the *mrpl40* phenotype, we treated wild-type embryos between 1 and 5 dpf with the mitochondrial ribosome inhibitor chloramphenicol (Cam)^25,26^ and tested their behavior at 6dpf (Figure 3A). Compared to embryo medium-treated larvae, Cam-treated larvae displayed a dose-dependent behavioral profile that was remarkably similar to *mrpl40* mutants (Figure 3B-3C). Specifically, we see that Chloramphenicol treatment recapitulates both Dark Flash (reduced movement metrics) and ASR (reduced sensitivity) phenotypes observed in *mrpl40* mutants (Figure 3B-3C). This result supports the hypothesis that *mrpl40* function in mitochondria is critical for distinct behaviors, including sensorimotor behaviors, in response to darkness and acoustic stimuli.

**Figure 3.**
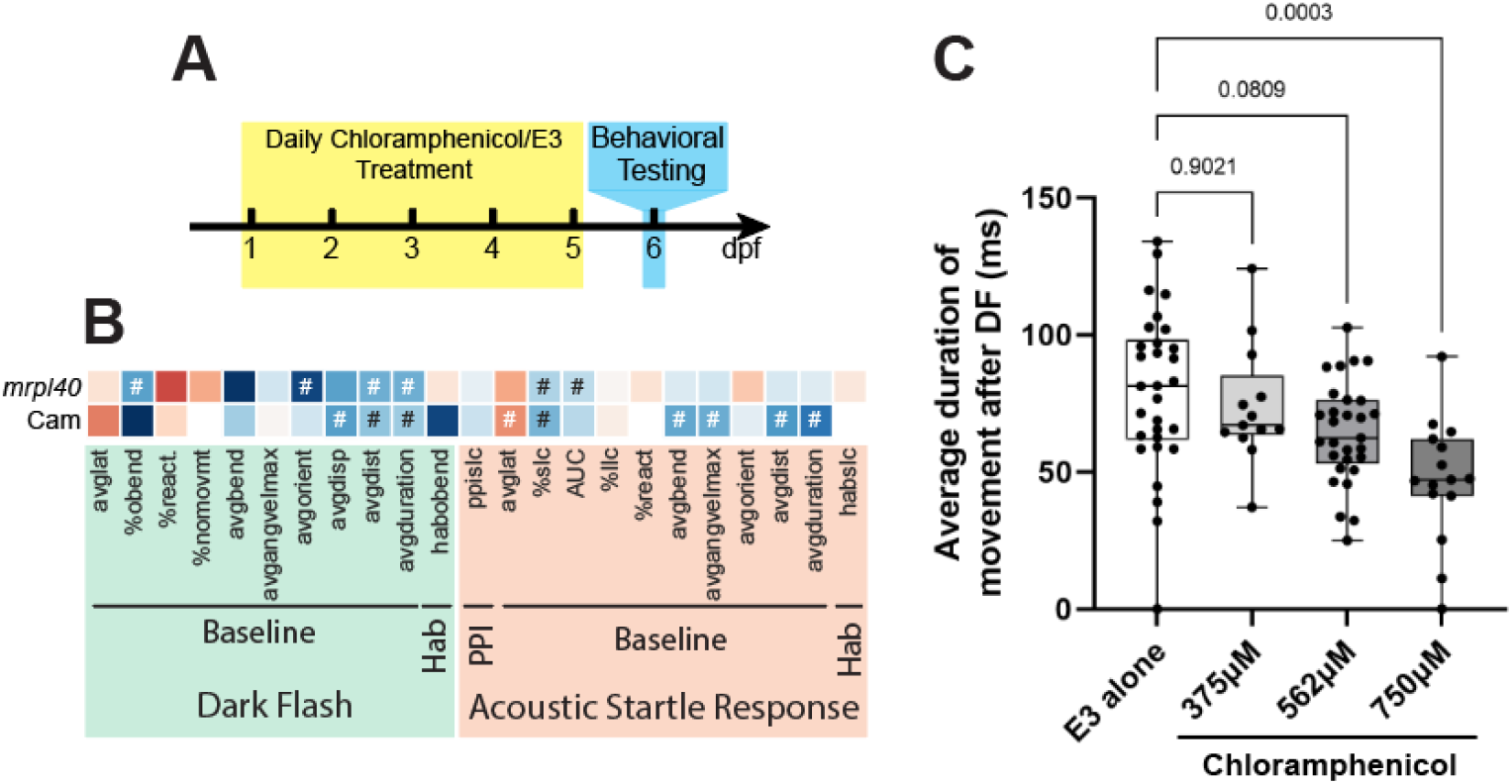
Pharmacologic inhibition of mitochondrial function phenocopies the *mrpl40* mutant phenotype. (A)Chloramphenicol experimental setup. Wild-type larvae were treated daily with chloramphenicol in E3 or with E3 alone and then behaviorally tested at 6dpf (B)Heatmap illustrating Chloramphenicol-treated (Cam) behavioral phenotype across DF and ASR behavioral metrics. *mrpl40* mutant phenotype heatmap is reproduced here from Figure 2 for comparison. Note the similarity between Cam-treated and *mrpl40* mutants across metrics assayed. Values that are significantly different from sibling/untreated controls based on Student’s t-test with a Bonferroni-corrected p-value of <0.05/23 are designated with #. See FFigure 2 legend for full description of heatmap. (C)Chloramphenicol-treated larvae have dose-dependent reductions in the average duration of movement following DF stimuli. One-way ANOVA, p-values are displayed on the plot.

### Mitochondrial disruption leads to alterations in brain volume

Individuals with 22q11.2DS exhibit alterations in brain size including a reduction in total brain volume and an increase in brain ventricular size^27,28^. At the same time, metabolic genes, including those involved in mitochondrial function, have been previously implicated in regulating cortical growth and size^29^. Thus, we examined whether disruption of mitochondrial functioning might cause behavioral phenotypes through an effect on brain volume. To determine if mitochondrial mutants and Cam-treated animals exhibit regional differences in brain volume we utilized deformation-based morphometry^30–32^ Briefly, larvae were raised to 6dpf, fixed, and stained for the brain structural marker total-ERK (tERK)^33^. Confocal stacks through the entire brain were then acquired on genotyped larvae and registered to a tERK reference brain^33^. Differences in deformation across the entire brain were computed for mutant/treated vs. wildtype/control larvae. Compared to controls, *prodha* mutant, *mrpl40* mutant, and Cam-treated (treated from 1-5dpf) larvae all displayed regional differences in brain volumes (Figures 4A-4C). Interestingly, differences appear similar across the two mutants and the mitochondrial translation inhibitor-treated larvae, with all three showing large reductions in telencephalon and mesencephalon regions and smaller increases in the rhombencephalon (Figures 4A-4E). *prodha* mutants and Cam-treated larvae also show additional reductions in the diencephalon (Figure 4D). We noted that reductions in brain volume were predominantly present in lateral brain regions, while increases tended to occur centrally with a distribution reminiscent of brain ventricles (Figures 4A-4C). To further assess this, using the Zebrafish Brain Browser^34^, we overlaid our volumetric data onto a registered confocal stack of *Tg(gfap:GFP)*, which labels populations of ventricular neural stem and progenitor cells (NSPCs). We found colocalization between increased brain volume signal and *Tg(gfap:gfp)* across all three conditions in multiple brain areas, suggesting possible increases in brain ventricular size and/or alteration of NSPC populations which reside along the ventricles. Importantly, compared to wildtype larvae, we observed reduced apoptosis in the brains of both *mrpl40* and *prodha* mutants (Figure S2), suggesting that rather than apoptosis, deficits in neurogenesis may underlie volumetric phenotypes.

**Figure 4.**
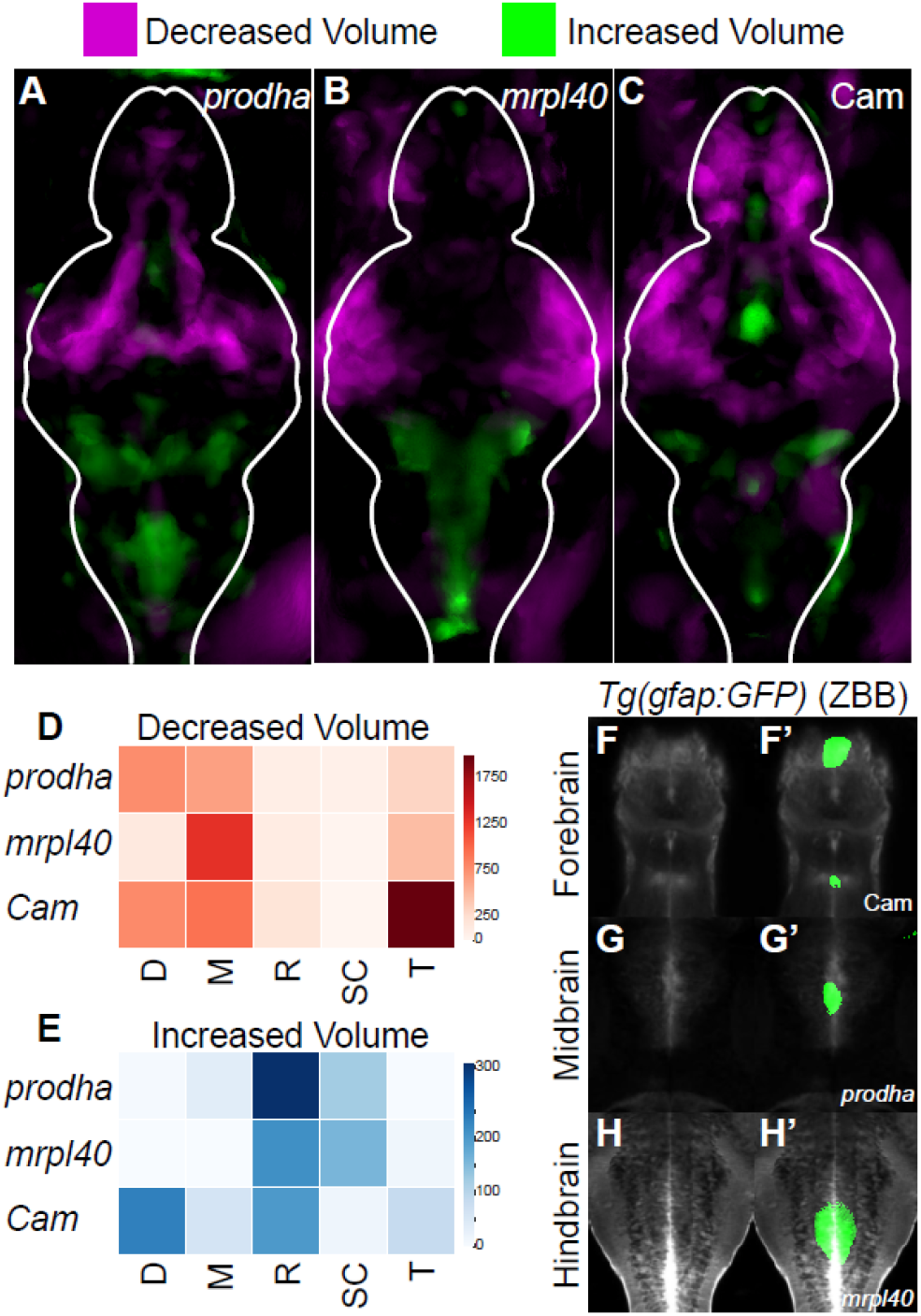
Mitochondrial disruption leads to alterations in brain volume. (A-C) Differences in brain volume in mitochondrial mutants and pharmacologically inhibited larvae. Deformation-based morphometry for *prodha* mutants (A), *mrpl40* mutants (B), and Chloramphenicol-treated (C) larvae shows differences in multiple brain areas. Images are sum projections of whole-brain z-stacks. Magenta designates areas that are reduced in volume compared to siblings/controls and Green designates areas that are increased in volume compared to siblings/controls. White outline indicates outline of brain. Analysis performed as in Thyme, *et al*.^32^. (D-E) Heatmaps illustrating degree to which volume signals in A-C occupy various brain regions in arbitrary units. Higher values indicate larger reductions (D)/increases (E) in volume in designated region. Values are scaled based on size of the region as performed in Randlett, *et al*.^33^. D, diencephalon; M, mesencephalon; R, rhombencephalon; SC, spinal cord; T, telencephalon. (F-H’) Colocalization of increases in brain volume with *gfap* reporter line. Average signal of a *Tg(gfap:GFP)* line that has been registered to the Zebrafish Brain Browser (ZBB) atlas was merged with increased volume signals for *prodha* mutants (G-G’), *mrpl40* mutants (H-H’), and Chloramphenicol-treated (F-F’) larvae. Increased volume colocalizes with GFP signal in multiple brain regions in the ventricle-occupying midline.

### *mrpl40* and *prodha* function independently from each other to regulate behavior

Given that both the proteins of *mrpl40* and *prodha* are localized to mitochondria and that mutants display similar behavioral and brain volumetric phenotypes, we wondered whether they regulate behavior through a common pathway. If so, double mutants are expected to display the same phenotypic severity when compared to single mutants. Conversely, if double mutants display a stronger phenotype, this would be consistent with the two genes acting in two separate pathways. To test this, we generated double mutants for *mrpl40* and *prodha* and analyzed their response to flashes of darkness (Figure 5). When presented with dark flash stimuli *mrpl40* mutants perform fewer O-bends than wildtypes, while *prodha* mutants perform O-bends at rates similar to wildtype (Figure 2 and 5). Analysis of *mrpl40* and *prodha* double mutants revealed a more severe phenotype than either single mutant alone (Figure 5), with 50% of animals not performing any O-bends (n=6/12 double mutants). This result indicates that while single mutants of *mrpl40* and *prodha* have similar behavioral and brain morphology phenotypes, they function in genetically distinct pathways, and that loss of one gene is at least partially compensated by the presence of the other gene.

**Figure 5.**
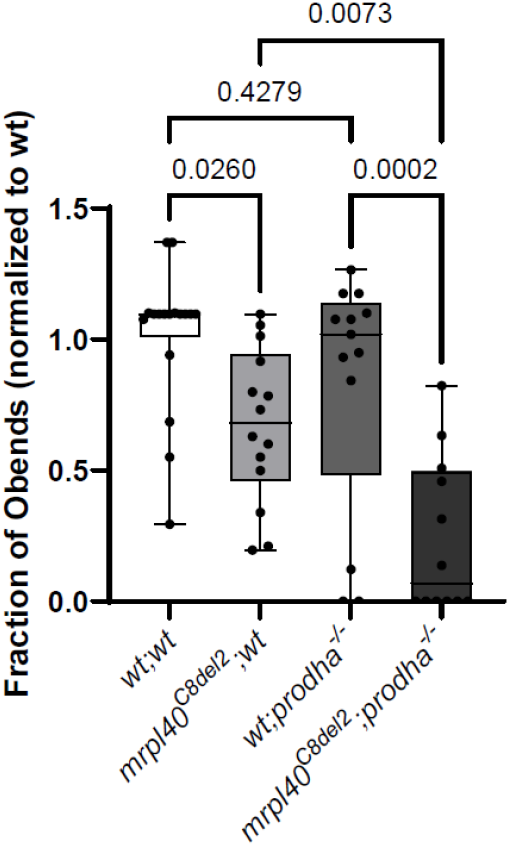
*mrpl40* and *prodha* function independently from each other to regulate behavior. *mrpl40* mutants (*mrpl40*^*C8del2*^*;wt*) perform fewer O-bends in response to dark flash stimuli. *prodha* mutants (*wt;prodha*^*-/-*^) have a trend to lower O-bends, but are not significantly different than wildtypes. *mrpl40* and *prodha double mutants (mrpl40*^*C8del2*^*;prodha*^*-/-*^) have a more severe phenotype than either single mutant. Note that this *mrpl40* mutation also disrupts gene *c22orf39*, though *c22orf39* single mutants do not have a phenotype. One-way ANOVA, p-values are displayed on the plot.

### *mrpl40* and *prodha* play distinct roles in NSPC proliferation

Volumetric analysis of *mrpl40* and *prodha* mutants revealed significant reductions in brain volume and abnormalities surrounding brain ventricles, where NSPCs reside. Recent studies have implicated mitochondria as important regulators of neurogenesis^16^, prompting us to consider the possibility that defective mitochondrial function in mutant animals may be linked to defects in NSPCs. To examine this possibility in more detail, we focused on the hindbrain, as both *mrpl40* and *prodha* mutants display significant volumetric changes in the hindbrain. Furthermore, the hindbrain contains the neuronal populations that regulate the ASR, which is disrupted in both mutants (Figure 2). Similarly, the hindbrain is also required for visuomotor behaviors that are disrupted in both mutants (Figure 2). At 5dpf, NSPCs marked by the multipotency marker Sox2 exist in two distinct populations in the hindbrain. First, NSPCs located in the dorsal hindbrain are highly proliferative, with the majority expressing the cell proliferation marker Proliferating cell nuclear antigen (PCNA) (Figure 6A, 7E). Second, NSPCs lining the ventral, central ventricular surface appear less proliferative, with only roughly 25% of Sox2+ cells expressing PCNA (Figure 6A, 6B, 6F). Unlike dorsal proliferative zone (PZ) NSPCs, NSPCs in the central PZ express both *gfap* and *fat4*, and likely represent radial glia-like cells (Figure S3).

**Figure 6.**
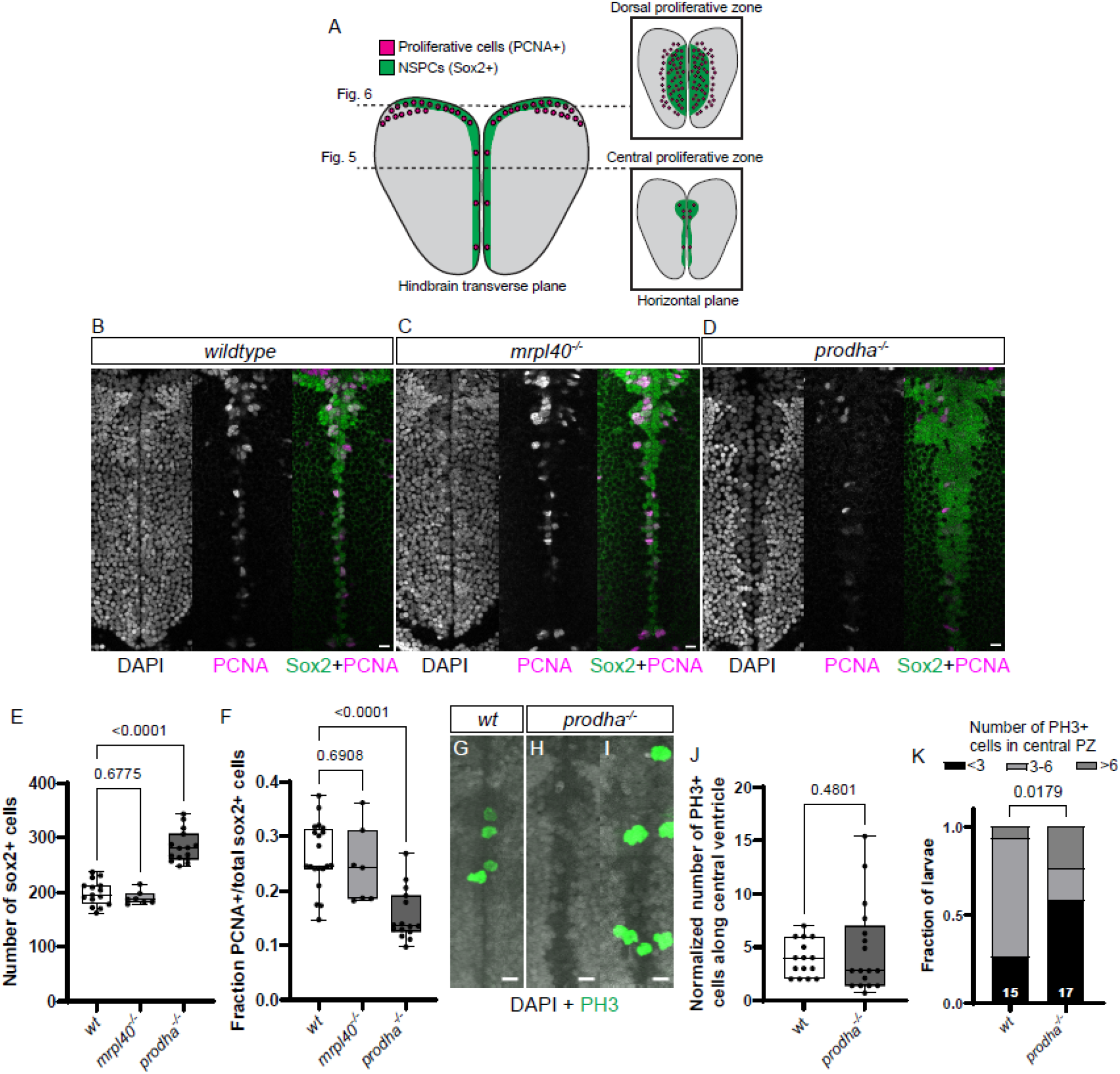
*prodha* mutant NSPCs are expanded and display abnormal proliferation in the central proliferative zone. (A) Diagram depicting NSPC populations in the zebrafish hindbrain at 5dpf. Transverse plane view shows highly proliferative NSPCs (Sox2+) and committed progenitors/neuroblasts (Sox-) located dorsally (Dorsal proliferative zone inset) and less proliferative NSPCs located centrally (Central proliferative zone inset). Insets are horizontal plane view. (B-F) Sox2 and PCNA staining in central proliferative zone in wildtypes (B), *mrpl40* mutants (C), and *prodha* mutants (D). Wildtypes and *mrpl40* mutants display similar patterns whereas *prodha* mutants have an expanded Sox2 domain and fewer PCNA positive cells. Images are single confocal slices. Scale bars 10μm. Quantification shows increase number of Sox2+ cells in *prodha* mutants (E), though a smaller fraction are PCNA+ (F). One-way ANOVA, p-values are displayed on the plots. (G-K) PH3 staining in central proliferative zone in wildtypes (G) and *prodha* mutants (H-I). *prodha* mutants show a similar average number of PH3+ cells as wildtypes (J), but the distribution of larvae is skewed (K). Unpaired two-way Student’s t-test for (I) and Chi-square for (K), p-values displayed on the plots. Images are maximum projections from confocal stacks. Scale bars 10μm.

We first assessed proliferation in the central PZ of *mrpl40* and *prodha* mutants by performing wholemount immunofluorescence with antibodies for the NSPC marker Sox2 and the proliferation marker PCNA (Figures 6B-6D). Wild-type larvae display Sox2+ NSPCs along the brain ventricular surface, and roughly a quarter of the NSPCs are PCNA+ (Figures 6B, 6E, 6F). In *mrpl40* mutants Sox2 and PCNA expression appeared indistinguishable from wild-type animals (Figures 6B, 6C, 6E, 6F). In contrast, *prodha* mutants displayed an expanded Sox2 population (Figures 6B, 6D). This expansion is consistent with our volumetric measurements (Figure 4A) that revealed increases in volume along the central hindbrain ventricle. Quantification revealed an increased number of Sox2+ cells in *prodha* mutants, though compared to wildtypes, the fraction of PCNA+ cells was decreased (Figures 6D-6F), suggesting fewer proliferative cells. In contrast, when we stained for the mitosis marker phospho-histone H3 (PH3), we failed to detect a difference in the average number of mitotic cells even after normalizing for the increase in Sox2+ cell numbers in *prodha* mutants (Figure 6J). However, *prodha* mutants displayed a greater variability in PH3+ cell numbers (Figures 6G-6I) and when we binned larvae into groups based on the number of PH3+ cells (Figure 6K), we found there were many more *prodha* mutants on each extreme of the range, indicating that *prodha* is required to regulate central PZ NSPC proliferation.

We next analyzed the dorsal PZ in *mrpl40* and *prodha* mutants. PCNA staining revealed that while the number of proliferative cells was similar between wildtype and *prodha* mutant hindbrains, there was a significant reduction in *mrpl40* mutants (Figures 7A-7D). In both mutants, however, DAPI nuclear staining revealed an enlarged ventricle (Figures 7A-7C), which may account, in part, for the increases in volume found in our volumetric experiments. To better understand the reduction in PCNA+ cells in *mrpl40* mutants and which cell types may be affected, we co-stained with Sox2 antibody. In wild-type animals, we observed Sox2+ cells located medially and expressing PCNA at high levels (PCNA^hi^). These PCNA^hi^/Sox2+ cells were abutted by weaker expressing PCNA cells (PCNA^lo^) laterally (Figure 7E). In contrast to PCNA^hi^ cells which also expressed Sox2, PCNA^lo^ cells did not express Sox2 and appeared to have smaller nuclei (Figures 7E). This is consistent with the presence of at least two populations of proliferative cells, with PCNA^hi^/Sox2+ cells likely representing highly proliferative uncommitted/intermediate progenitors and PCNA^lo^/Sox2-cells representing more committed progenitors or neuroblasts. While the number of PCNA^hi^/Sox2+ cells in *mrpl40* mutants was unaffected (Figures 7E-7G) when compared to wild-type animals, PCNA^hi^/Sox2+ cells displayed abnormal morphology, appearing elongated (Figures 7H-7I). In contrast to PCNA^hi^/Sox2+ wild-type cells, in *mrpl40* mutants, the number of PCNA^lo^/Sox2-cells was greatly reduced (Figures 7E-7G). We wondered whether a proliferative defect in PCNA^hi^/Sox2+ cells may be underlying the reduction in PCNA^lo^/Sox2-cell numbers. To test this idea, we analyzed the spatial distribution of PH3 expression within the dorsal PZ. As PCNA^lo^/Sox2-cells are located laterally, we reasoned that PH3 expression in cells located centrally along the ventricle would represent mitotic PCNA^hi^/Sox2+, whereas PH3+ cells located more laterally would be a mixture of PCNA^hi^/Sox2+ and PCNA^lo^/Sox2-cells. We counted PH3+ cells within the dorsal PZ and grouped them based on whether the cells were located in the first cell layer adjacent to the ventricle or in more lateral regions. Consistent with the overall decrease in dorsal PZ PCNA+ cells, we observed a reduction in lateral PH3+ cells in *mrpl40* mutants (Figures 7J-7K). Moreover, we also observed a reduction in central PH3+ cells, suggesting that reduced proliferation of PCNA^hi^/Sox2+ cells likely contributes to the observed reduction in PCNA^lo^/Sox2-cell numbers (Figures 7J-7K). Together, our results indicate that both *prodha* and *mrpl40* are required for normal NSPC proliferation, though they appear to regulate distinct populations of NSPCs: *prodha* affects proliferation in the central PZ, whereas *mrpl40* affects proliferation in the dorsal PZ. Furthermore, loss of either gene leads to similar brain morphology and behavioral outcomes and loss of both genes aggravates behavioral phenotypes. Taken together, our data reveal a previously unappreciated link between 22q11.2DS-relevant behavioral phenotypes, mitochondrial function, and neurogenesis.

**Figure 7.**
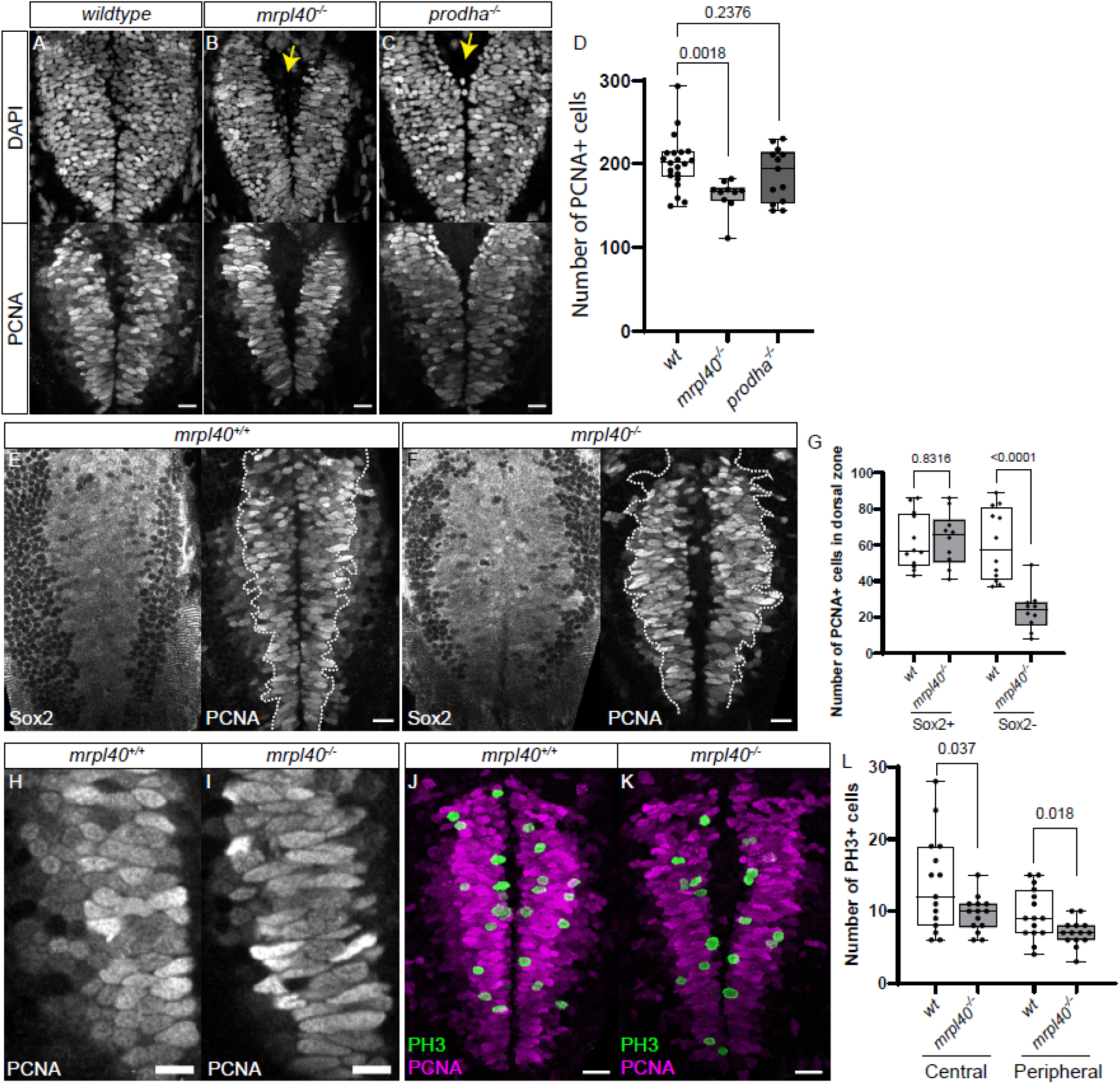
*mrpl40* mutants have fewer NSPCs and a reduction in NSPC proliferation in the dorsal proliferative zone. (A-D) PCNA and DAPI antibody staining in wildtypes (A), *mrpl40* mutants (B), and *prodha* mutants (C) in dorsal proliferative zone. *mrpl40* mutants have fewer PCNA+ cells than wildtypes. *prodha* mutants have similar numbers to wildtypes. Quantification in (D). One-way ANOVA, p-values are displayed on the plots. Both *mrpl40* and *prodha* mutants appear to have increased ventricle size (yellow arrows). Images are single confocal slices. Scale bars 15μm. (E-G) Sox2/PCNA antibody staining in wildtypes (E) and *mrpl40* mutants (F) in dorsal proliferative zone. *mrpl40* mutants show a reduction in the number of Sox2-/PCNA+ cells but no difference in the number of Sox2+/PCNA+ cells (G). Quantification in (G). Unpaired two-way Student’s t-test, p-values displayed on the plot. Images are single confocal slices. Dotted line indicates boundary of Sox2+ cells. Scale bars 15μm. (H-I) Proliferative cells in dorsal hindbrain of *mrpl40* mutants (I) display elongated nuclei compared to wildtypes (H). Images are single confocal slices. Scale bars 10μm. (J-L) PH3/PCNA antibody staining in wildtypes (J) and *mrpl40* mutants (K) in dorsal proliferative zone. *mrpl40* mutants show a reduction in the number of PH3+ cells located both centrally along ventricle and more peripherally. Quantification in (L). Unpaired two-way Student’s t-tests, p-values displayed on the plot. Images are maximum projections from confocal stacks. Scale bars 20μm.

## Discussion

22q11.2DS predisposes to multiple NDDs and represents one of the greatest risk factors for schizophrenia, yet it has been difficult to define which genes within the region contribute to brain and behavioral phenotypes. Part of this difficulty is likely due to the idea that rather than single genes acting alone to impart disease risk, multiple genes within the deleted region act together through shared biological pathways/functions to drive pathology. However, the identity of these shared pathways and the identity of the relevant genes has remained unclear. Here, we identify previously unappreciated roles for the *mrpl40* and *prodha* genes in NSPC proliferation, and provide compelling evidence that both genes function in genetically separate pathways to regulate mitochondrial-supported NSPC proliferation. Furthermore, while our results support previous studies implicating mitochondrial function in 22q11.2DS pathology, they indicate that defects in NPSCs may precede post-mitotic defects. Finally, our results provide the first functional link between mitochondrial gene dysfunction, brain volumetric changes, and 22q11.2DS-relevant behavioral phenotypes. Thus, our results strongly support the notion that mitochondrial dysfunction plays an early and central role in the etiology of neurodevelopmental disorders associated with 22q11.2DS.

Previous studies have suggested that mitochondrial dysfunction may be involved in 22q11.2DS pathogenesis. Indeed, at least six of the genes in the deleted region have been shown to localize to mitochondria^10^, with several displaying behavioral phenotypes in mouse knockouts^14,15,35,36^. Further, iPSC-derived neurons from patients with the deletion display reduced levels of electron transport chain components and ATP^11,12^. Despite evidence that suggests NSPC proliferation may be defective in 22q11.2DS^3,37,38^, studies to date have focused on mitochondrial roles in post-mitotic neurons. Our data indicates that two evolutionary conserved mitochondrial genes within the 22q11.2 deleted region are required for zebrafish NSPC function, implicating NSPC mitochondrial dysfunction as a potential mechanism in 22q11.2DS pathology. We hypothesize that mitochondrial dysfunction resulting from cumulative effects of deleterious mutations in multiple genes in the region results in NSPC deficits. Indeed, in addition to our findings regarding *mrpl40* and *prodha*, a recent study demonstrated that *mrpl40* and another mitochondrial gene in the deleted region, *slc25a1*, biochemically interact to maintain mitochondrial ribosomal integrity^39^. Our hypothesis is further supported by an increasing recognition of mitochondria as key regulators of NSPC proliferation and neurogenesis^16,40^ and recent data suggesting that mitochondrial dysfunction may serve as a general pathological mechanism underlying multiple NDDs^41,42^. Therefore, understanding the roles that mitochondria serve in NSPCs is crucial to furthering our understanding of 22q11.2DS and NDDs in general.

Mitochondrial ribosomal protein L40 (*mrpl40*) is a component of the mitochondrial ribosome that is required for translation of mitochondrially encoded genes, many of which are involved in oxidative phosphorylation (OxPhos). Previously, *mrpl40* has only been examined in the context of post-mitotic neuron function. Heterozygous loss of *mrpl40* in forebrain-like excitatory neurons in culture leads to reductions in OxPhos complexes and ATP levels that phenocopy deficits observed in 22q11.2DS patient-derived neurons^11^. Heterozygous *mrpl40* mouse knockouts display deficits in working memory that have been attributed to abnormal intracellular calcium buffering^43^. Knockdown of *mrpl40* in *Drosophila* leads to abnormal larval neuromuscular junction synapse development and function and sleep phenotypes^39^. Here we show a previously unrecognized role for *mrpl40* in regulating brain size and NSPC proliferation. Due to its role in OxPhos gene translation, we hypothesize that mitochondrial OxPhos genes are required for NSPC proliferation and maintenance of brain volume. This is supported by our mitochondrial translation inhibition experiments that revealed behavioral and brain volume changes similar to those in *mrpl40* mutants. This would also be consistent with data from *Drosophila*^40^ and mouse^46,47^ studies that found that when OxPhos is disrupted in different contexts, NSPC proliferation is reduced. Interestingly, however, we find that different populations of NSPCs exhibit differential sensitivity to the loss of *mrpl40* function. Specifically, *mrpl40* mutants display reduced NSPC proliferation in the dorsal PZ of the hindbrain, while radial glia-like cells in the central PZ appear unaffected. One possible explanation for this is that NSPC populations have different reliance on mitochondria. Indeed, multipotent neural stem cells are mainly glycolytic and shift to relying on mitochondria as intermediate progenitors and during commitment^48,49^. Therefore, it is possible that the radial glia-like central PZ NSPCs are more reliant on glycolysis whereas dorsal PZ NSPCs require mitochondria. Another possibility may relate to differences in proliferation between these populations. Specifically, the more highly proliferative nature of dorsal PZ NSPCs may mean that they are more reliant on mitochondria. Interestingly, OxPhos disruption in other contexts has been shown to alter cell cycle dynamics^44,45^, so it is possible that highly proliferative cells may be more vulnerable to mitochondrial dysfunction. Understanding the downstream results of loss of *mrpl40* function and the specificity of the NSPC proliferative phenotypes are important questions that remain unanswered.

Proline dehydrogenase (*prodh*) is localized to mitochondria and is responsible for the first step of L-proline metabolism. Humans with homozygous mutations in *prodh* display multiple NDDs including developmental delay, ID, ASD, and SZ^50^, as well as elevated levels of L-proline. To date, studies of *prodh*’s role in brain function and behavior have mainly focused on the role of hyperprolinemia in post-natal post-mitotic neurons^13,14^. However, studies do not support a link between L-proline levels and clinical phenotype^50^, suggesting roles outside of maintaining L-proline levels that are likely involved. Our results indicate that *prodh* has an important role in NSPC development, which is more in line with observed NDD clinical phenotypes. Specifically, we find that *prodha* mutants display a reduction in brain size and an expanded central PZ with alterations in NSPC proliferation. Although its role in NSPC development has not been previously examined, *prodh* has been extensively investigated in other highly proliferative cells, such as cancer cells where it can act as either a tumor suppressor or oncogene depending on context^51^. The mechanisms underlying the NSPC phenotypes we observed are unknown, though there appear to be several possibilities that warrant further exploration. L-Proline oxidation via *prodh* leads to reducing equivalents of NADH/FADH_2_ that can feed OxPhos to serve NSPC energetic needs. This scenario seems less likely since *mrpl40* mutants display a different NSPC phenotype. Alternatively, *prodh* has been shown to promote reactive oxygen species (ROS) formation in several contexts^52–54^ and low levels of exogenous ROS have been shown to promote proliferation of NSPCs^55^ and other stem cells^56^. L-Proline’s metabolism also leads to glutamate, which can contribute to the TCA cycle and play critical roles for forming other cellular substates important for highly proliferative cells. Furthermore, the TCA cycle has recently been identified as a key hub of cell fate decisions in embryonic stem cells^57^, so it is possible that TCA substrates may perform more instructive roles during neurogenesis. Further experiments will need to be performed to identify the precise mechanism underlying *prodha*’s role in NSPCs. It is important to note that the *prodha* NSPC phenotype appears to be more complex than the *mrpl40* phenotype. While the fraction of PCNA+ central PZ NSPC cells is decreased, PH3 staining reveals that there is an increase in both the fraction of larvae that show decreased mitoses and the fraction of larvae that have increased mitoses. One possible explanantion is that *prodha* functions to regulate mitoses and that its loss leads to under or over proliferation depending on cellular context. Alternatively, loss of *prodha* might first result in NSPC hyperproliferation and ultimately reduced proliferation over time, which could potentially explain the increase in central PZ NSPC numbers observed in *prodha* mutants. Further studies will be required to dissect *prodha*’s complex role in NSPC biology.

Together, our data presents NSPC proliferation defects as a potential core component of 22q11.2DS pathology. Our work complements previous studies that have shown that two other genes in the deleted region, *dgcr8*^58,59^ and *ranbp*^60^, also affect NSPC proliferation. However, it remains to be determined to what extent NSPC proliferation deficits lead to behavioral outcomes and through which mechanisms. To fully appreciate how mitochondrial dysfunction caused by loss of *mrpl40* and *prodha* may lead to behavioral phenotypes will require careful examination of progenitor and post-mitotic cell diversity.

## Supporting information

Primary Supplemental Data

Supplemental Zip file 1 - python tracking scripts

## Acknowledgements

The authors would like to acknowledge the University of Pennsylvania Cell and Developmental Biology Microscopy Core and the Genomic Analysis Core DNA Sequencing Facility and the University of Alabama at Birmingham Research Computing team. We also thank the Granato lab members for feedback on experimental design and the manuscript. This work was supported by grants to M.G. (NIH R01 NS118921) and P.D.C. (NIH R25 Pilot funding). P.D.C. was supported by T32MH019112 and R25MH119043. S.T. was supported by R00MH110603.

## Author contributions

Conceptualization, P.D.C. and M.G.; Investigation, P.D.C. and I.L.; Formal Analysis, P.D.C. and S.T.; Resources and Supervision, M.G.; Funding Acquisition, P.D.C. and M.G.; Writing – Original Draft, P.D.C.; Writing – Review and Editing, P.D.C., S.T. and M.G.

## Methods

### RESOURCE AVAILABILITY

#### Lead Contact

Michael Granato

#### Materials availability

Further information and requests for resources and reagents should be directed to and will be fulfilled by the lead contact, Michael Granato

#### Data and code availability

All data reported in this paper will be shared by the lead contact upon request. All original code is available in this paper’s supplemental information. Any additional information required to reanalyze the data reported in this paper is available from the lead contact upon request.

### EXPERIMENTAL MODEL AND SUBJECT DETAILS

Experiments were conducted on 5-6 dpf larval zebrafish (Danio rerio, TLF strain) raised in E3 medium at 29°C on a 14:10 hr light cycle. At this developmental stage the sex of the organism is not yet determined. Breeding adult zebrafish were maintained at 28°C on a 14:10 hr light cycle. Mutant alleles were generated using CRISPR-Cas9 mutagenesis. For single gene mutants, two gRNAs flanking a known functional domain or highly conserved region were designed using ChopChop v2. Alt-R CRISPR-Cas9 crRNAs (IDT) were annealed with tracrRNA (IDT), complexed with Cas9 protein (PNA Bio), and injected into 1-cell stage embryos. F_0_ injected larvae were raised and outcrossed to identify and establish heterozygous carrier lines. For combination gene mutants, four gRNAs were used: two gRNAs targeting the gene flanking one end of the deletion were injected together with two gRNAs targeting the gene flanking the other end in order to delete the intervening region. Mutant lines were identified and subsequently genotyped by PCR, using primers flanking the gRNA target sites. Allele sequences were obtained by Sanger-sequencing of PCR products. For a subset of alleles, genotyping was also performed with the KASP method with proprietary primer sequences (LGC Genomics). This method was validated using the PCR method. All animal protocols were approved by the University of Pennsylvania Institutional Animal Care and Use Committee (IACUC).

### METHOD DETAILS

#### Behavior recording

Behavior experiments were performed on 6dpf larvae. For each behavioral run, approximately 50 larvae were used for each mutant allele. Larvae were arrayed in 100-well plates fabricated from 2mm thick clear acrylic sheets (McMaster-Carr). Wells were 9mm in diameter and 2mm in depth and arrayed 10x10. The plate was illuminated from below with IR-850nm LED arrays (IR100, CMVision) which were diffused using a white acrylic sheet. Images were recorded from above with 1024x1024 resolution using a Photron UX50 camera, Quantaray 50mm 1: 2.8D Macro for Nikon AF lens, and IR long-pass filter (LP800-55, Midwest optical). To provide acoustic vibrational stimuli (2ms duration, 1000 Hz waveforms), the plate was attached via an aluminum rod to a vibrational exciter (4810; Brüel and Kjaer, Norcross, GA). White light (MCWHL2, Thorlabs) was provided from above (700mW, 6500K). Video recording, white light LED, and vibrational stimuli were triggered by a NI PCI-6221 M Series DAQ^23^.

The full behavioral paradigm consisted of the following described blocks in the stated order. VMR: Lights OFF (10mins, unrecorded) > Lights ON (8mins, 20fps) > Lights OFF (8mins, 20fps) > Video saved (∼12mins). LF: 15 stimuli of white light of 500ms duration with 30s interstimulus interval (ISI). Recording 500ms post-stimulus at 500fps (8mins) > Video saved (∼2mins). DF: Lights ON (5mins, unrecorded) > 14 stimuli of darkness of 1s duration with 30s ISI (recorded) > 14 stimuli of darkness of 1s duration with 10s ISI (unrecorded) > 14 stimuli of darkness of 1s duration with 10s ISI (recorded) > 14 stimuli of darkness of 1s duration with 10s ISI (unrecorded) > 14 stimuli of darkness of 1s duration with 10s ISI (recorded). Recording 1s post-stimulus at 500fps (∼23mins total) > Video saved (∼12mins). ASR: 10 stimuli of low intensity with 20s ISI > 10 stimuli of medium intensity with 20s ISI > 10 repetitions of the following with 20s ISI PP1-PPI2-PPI3-PPI4-High Intensity (where PPI1: low intensity prepulse, 50ms prior high intensity, PPI2: low intensity prepulse, 300ms prior high intensity, PPI3: medium intensity prepulse, 50ms prior high intensity, PPI4: medium intensity prepulse, 300ms prior high intensity, High Intensity: no prepulse) > 30 stimuli of high intensity with 1ISI. Recording 200ms post-stimulus at 500fps (∼24mins total) > Video saved (∼6mins). The behavioral paradigm took place over an approximately two-hour period during the daytime hours from 8AM to 4PM. Following behavioral experiments, larvae were individually genotyped as described above.

#### Larval tracking

Offline tracking of recorded videos was performed with Python. Each image was background subtracted, Gaussian blurred, thresholded, and the centroid identified to identify larvae. For LF, DF and ASR assays five points along the tail were also identified as previously described^61^. The orientation of the larvae was determined as the vector between the first point along the body and the centroid and the curvature was defined as the sum of the angles between all of the points along the larva.

#### Behavioral metrics

A total of 94 metrics were computed for each behavioral run and they are described below.

*VMR (60):* Number of bouts, Total distance travelled (pix), Total time moved (ms), Average speed (pix/ms), Average distance from center of the well (pix), Fraction of time spent in out rim of well, Average distance/bout (pix), average displacement/bout (pix), Average time/bout (ms), Average speed/bout (pix/ms). A bout was defined as the displacement of the centroid crossing a certain threshold (0.7 pixels) for at least two consecutive frames. For a bout to end, two consecutive frames had to have a displacement below this threshold. Centroid movements were only considered if they occurred during periods defined as bouts. Behavioral metrics were considered for the first and last minutes of the Light ON and Light OFF segments and for the entirely of each of the Light ON and Light OFF segments for a total of six bins for each metric, and a total of 60 metrics for the VMR assay

*LF (9):* Frequency of reacting, frequency of no movement, Average latency to movement (ms), Average maximum bend angle (rad), Average maximum angular velocity (rad/frame), Average change in orientation (rad), Average displacement (pix), Average distance (pix), Average duration (ms). Larvae were classified as reacting if their maximum curvature was > 0.5rad and latency of movement was not too early (>15ms).

*DF (11):* In addition to the metrics for LF, Frequency of O-bend response and Habituation of the O-bend response were calculated. Larvae were classified as performing an O-bend if their maximum curvature was >1.75rad or reacting if it was > 0.5rad but less than 1.75rad as long as latency of movement was not too early (>15ms). O-bend habituation was calculated as (1- [Obend frequency for last 14 stimuli]) ÷ [Obend frequency for first 14 stimuli].

*ASR (14):* In addition to the metrics for LF, Frequency of SLC response, Frequency of LLC response, Sensitivity Index of the ASR (AU), Pre-pulse inhibition of the ASR, and Habituation of the ASR were calculated. Larvae were classified as performing an SLC if their maximum curvature was >0.8rad and the latency was <20ms, performing an LLC is their maximum curvature was >0.8rad and the latency was >20ms, or reacting if it was > 0.5rad but less than 0.8rad. ASR habituation was calculated as (1-[SLC frequency for last 10 High intensity stimuli with 1s ISI]) ÷ [SLC frequency for 10 High intensity stimuli with 20s ISI]. ASR PPI was calculated as (1-[SLC frequency for 10 PPI4 runs]) ÷ [SLC frequency for 10 High intensity stimuli with 20s ISI]. ASR Sensitivity index was calculated by measuring the area under the curve of SLC frequency versus stimulus intensity.

#### Chloramphenicol treatment

Wild-type TLF larvae were raised in E3 medium, dechorionated, and treated with either Chloramphenicol (C0328, Sigma) dissolved in E3 or E3 medium alone starting at 1dpf. Media was replaced daily up until 6dpf when behavioral experiments were performed. Behavior experiments were performed using serial dilutions of Chloramphenicol to show dose-response (375-750μM), whereas dose used for brain volumetric analysis was 562μM.

#### Deformation-based morphometry

Deformation-based morphometry was performed as described previously^32^. Briefly, whole-brain confocal stacks were registered to the previously reported 6-dpf *nacre* mutant (*mifta*^*− /−*^) larvae stained with anti-total ERK using CMTK (http://www.nitrc.org/projects/cmtk/) via the Fiji-CMTK-registration-runner-GUI (https://github.com/sandorbx/Legacy-Fiji-CMTK-registration-runner-GUI) with the following parameters: Threads 8, Initial exploration step 52, Coarsest resampling 8, Refine grid 3, Grid size 80, Accuracy 1. The jacobian was calculated using reformatx in CMTK, followed by smoothing the images with a custom Fiji script PrepareJacobianStacksForMAPMapping_cluster.m as previously described^32^ (github repository: sthyme/ZFSchizophrenia/cluster_pErk_imageprocessing/). To compare volumetric stacks with *gfap* expression, the *Tg(gfap:gfp)* reference stack was obtained from the Zebrafish Brain Browser (http://vis.arc.vt.edu/projects/zbb/) and merged with volumetric stacks using Fiji.

#### Immunohistochemistry

Larvae were raised to 5-6dpf at which point they were fixed in 4% paraformaldehyde in PBS. Larvae were stained as previously described^33^. Briefly, larvae were washed with PBS+0.25% Triton X-100 (PBT) and pigment was removed using a 1.5% H_2_O_2_/0.5% KOH solution at room temperature (RT) for 15mins followed by further PBT washes. Antigen recovery was performed with 150 mM Tris-HCl, pH 9 for 5 minutes at RT followed by 15 minutes at 70°C, followed by PBT washes. Permeabilization was performed with 0.05% Trypsin-EDTA for 5 minutes on ice followed by PBT washes. Larvae were then blocked with PBT+1%bovine serum albumin+2% normal goat serum (NGS)+1% DMSO for 1 hour at RT followed by overnight incubation at 4°C with primary antibodies and DAPI in blocking solution without NGS. The following day PBT washes were performed followed by overnight incubation at 4°C with secondary in blocking solution without NGS. Following further PBT washes, larval tails were removed with a razor blade and used for genotyping. Following genotyping, larvae were mounted in 1.5% low melting-point agarose in PBS dorsal side down in a glass-bottomed dish and imaged with an inverted confocal microscope (Zeiss LSM880) using either a 20X air objective for whole brain analysis, or a 40X water objective for hindbrain analysis. Whole-brain images were acquired in two stacks at a voxel size of ∼0.8×0.8×2 µm (x×y×z) which were then stitched together using Zeiss Zen online stitching. Hindbrain images were acquired in 2-3 stacks at a voxel size of ∼0.8×0.8×2 µm (x×y×z) to cover most of the brain and were then stitched together using Zeiss Zen online stitching. Primary antibody concentrations used were: Mouse Anti-PCNA Antibody (1:500), Rabbit Anti-active Caspase-3 (1:500), Rabbit Anti-Sox2 (1:200), Rabbit Anti-phospho-Histone H3 (1:500), Mouse Anti-total-ERK (1:500). Alexa Fluor secondary antibodies were used at 1:500 dilution.

### QUANTIFICATION AND STATISTICAL ANALYSIS

#### Behavioral quantification

Behaviors were quantified, analyzed, and visualized with Python. Approximately 50 larvae from a heterozygous in-cross for each mutant allele were run through the behavior paradigm. Following genotyping, mutant larvae were compared to sibling larvae across all 94 behavioral metrics with two-tailed, unpaired Student’s t-tests. If any behavioral metric had p<0.05, an additional independent experiment was performed (expect for *cdc45* where only a single experiment was performed for technical reasons). In this independent experiment, p values were compared with the first experiment to identify if any behavioral metrics were in common. This was the case for 13 mutant lines and they had varying numbers of metrics in common ranging from 1 to 53. Those mutants that only had differences in metrics in the VMR assay and which only occurred within a single 1-minute bin of time and in no others during the 16 minute assay (*slc25a1b* (1 metric) and *scarf2* (4 metrics)) were felt to less likely represent true differences and were not considered further. To account for multiple comparisons, the data for both experiments was then pooled and subjected to further statistical stringency using a Bonferroni correction of p=0.05/94. Prior to pooling, to account for run-to-run variation in behavior, behavioral metrics were first normalized within their experiment by dividing all values by the average sibling value. Normalized values were then pooled and statistics performed. Nine mutants had behavioral metrics that survived correction. One of the nine mutants (*dgcr2*), was significant only for habituation of the SLC response in the ASR assay. However, due to technical limitations and loss of the line, the ASR assay was only performed once. Since habituation of the SLC was only significant for the last 10 stimuli of the habituation paradigm and not the first 10 or the middle 10 stimuli, this was deemed less likely to be a true difference. The eight remaining mutants had a range of 5 to 47 behavioral metrics below the corrected p value. These are the data represented by the heatmaps shown in Figures 2G and S1.

#### Deformation-based morphometry

Deformation-based morphometry was analyzed as described previously^32^. The Mann-Whitney U statistic Z score is calculated for each voxel, comparing between the mutant/treated groups and control groups. The significance threshold was set based on a false discovery rate (FDR) where 0.005% of control pixels would be called as significant. Following this, the extent to which volumetric changes overlapped within different Z-Brain atlas defined regions was performed by processing the volumetric stacks with the previously reported “ZBrainAnalysisOfMAPMaps.m” Matlab function^33^.

#### Immunohistochemistry quantification

All image visualization and processing was performed with Fiji. *Activated Caspase 3:* Positive cells were manually counted by manually progressing through stacks. The forebrain, tectum, and hindbrain were defined by anatomical landmarks visualized with DAPI co-staining. *Sox2/PCNA in central PZ:* For Figures 6B-6D, a single confocal slice at a comparable anatomical location ∼50μm below dorsal surface of hindbrain was used for analysis. DAPI signal was gaussian blurred, maximum filtered with radius of 2pixels, unsharp masked with radius 5 pixels and mask weight 0.9, thresholded, and then watershed function used. Sox2 signal was then thresholded and used to mask DAPI image, to leave only Sox2+ nuclei which were subsequently counted with the Analyze Particles function. PCNA signal was then gaussian blurred, thresholded, and used as a mask on Sox2+ nuclei, to leave Sox2+/PCNA+ nuclei which were subsequently counted with the Analyze Particles function. *PH3 in central PZ:* For Figures 6G-6K, PH3+ cells were manually counted by manually progressing through a 100μm thick confocal stacks. DAPI/PCNA co-staining allowed visualization of the central ventricle. Number of PH3+ cells in *prodha* mutants was normalized by dividing the observed number of PH3+ cells by the average fractional increase of Sox2+ cells observed in *prodha* mutants compared to wildtypes. *Sox2/PCNA in dorsal PZ:* For Figures 7A-7D, a single confocal slice at a comparable anatomical location ∼10μm below dorsal surface of hindbrain was used for analysis. DAPI signal was gaussian blurred, maximum filtered with radius of 2pixels, unsharp masked with radius 5 pixels and mask weight 0.9, thresholded, and then watershed function used. PCNA signal was then thresholded and used to mask DAPI image, to leave only PCNA+ nuclei which were subsequently counted with the Analyze Particles function. For Figures 7E-7G, Sox2+/PCNA+ and Sox2-/PCNA+ cells were manually counted in an ROI encapsulating a 93µmx122µm segment of one of the lateral hindbrain PCNA+ segments in a single z-slice ∼10µm below the dorsal most aspect of the hindbrain. *Hindbrain PH3:* Positive cells were manually counted by manually progressing through stacks. DAPI co-staining allowed for distinction of those that were within the first cell layer along the ventricles vs. those that were located more peripherally. One-way ANOVA, Chi-square, or two-tailed, unpaired Student’s t-tests were used for immunohistochemistry statistics as indicated in Figure legends.

## ADDITIONAL SUPPLEMENTAL INFORMATION

**Supplemental Table 1** (excel file) – Single gene mutations, gRNA sequences, and genotyping primer sequences

**Supplemental Table 4** (excel file) – Combination gene mutations, gRNA sequences, and genotyping primer sequences

**Supplemental Table 5** (excel file) - Annotated example of behavioral output for single behavioral example.

**Supplemental Table 6** (excel file) – Summary data for eight mutant lines with behavioral phenotypes. Two independent experiments performed for ech line. Table denotes behavioral metrics where p<0.05 in both experiments.

**Supplemental Table 7** (excel file) – Normalized and pooled data for each of the eight mutant lines with behavioral phenotypes. p values also included for each behavioral metric.

**Supplemental Zip file 1** – Python scripts used for behavioral tracking.

