## Supplementary material for "Mitochondrial genes in the 22q11.2 deleted region regulate neural stem and progenitor cell proliferation": Primary Supplemental Data

**Supplemental Figure 1. Single gene mutants are phenocopied by second allele disrupting same gene**

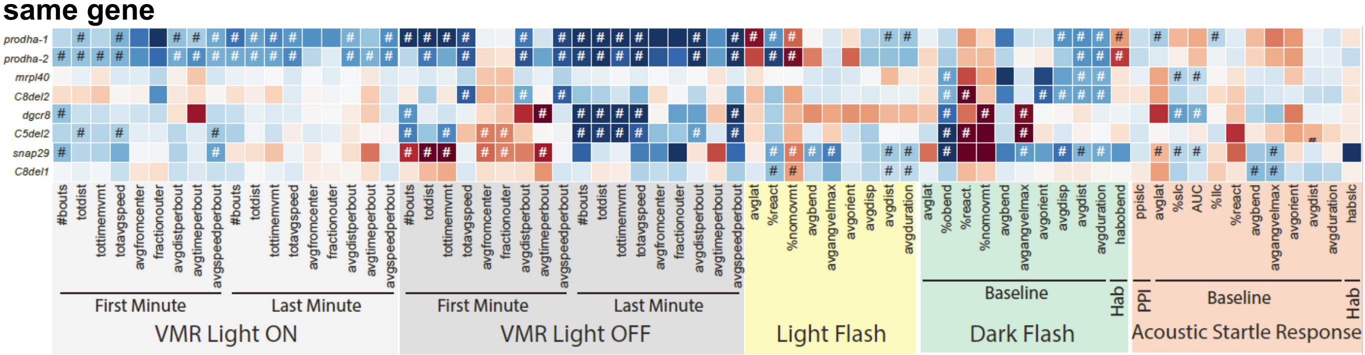


Heatmap illustrating mutant phenotypes across behavioral metrics. Each row is an individual mutant line and all columns represent a behavioral metric. Color of each box represents difference of average mutant value compared to siblings expressed as number of standard deviations (SD). Blue boxes indicate the value is lower in the mutant and Red boxes indicate the value is higher in the mutant. Mutant values that are significantly different from siblings based on Student's t-test with a Bonferroni-corrected p-value of 0.05/94 are designed with #. VMR behavioral metrics are highlighted in gray, LF in yellow, DF in green, and ASR in orange. Two independent single gene *prodha* mutant lines display the same phenotype across behavioral metrics. Combination gene lines *C8del2*, *C5del2*, and *C8del1* that overlap *mrpl40*, *dgcr8*, and *snap29* respectively, phenocopy single gene lines, indicating single genes are the drivers of the combination line phenotypes.

**Supplemental Figure 2. *mrpl40* and *prodha* mutants do not display increased apoptosis**

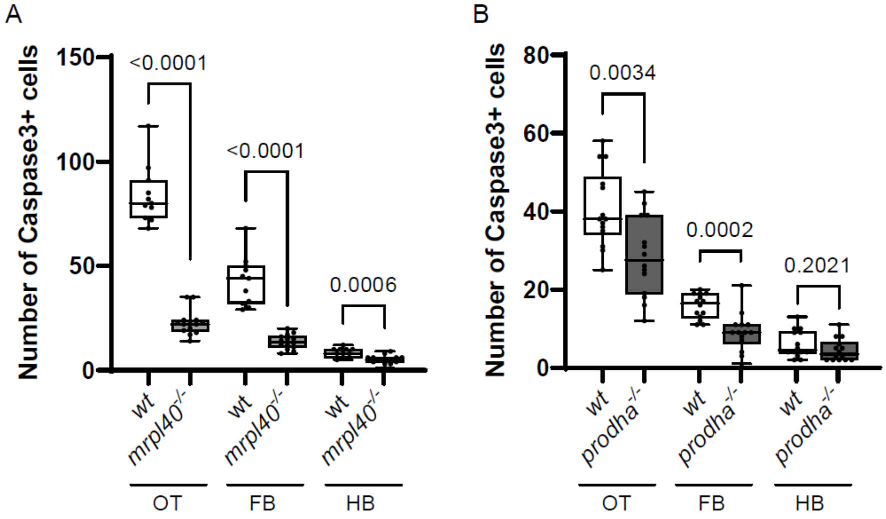


(A) Caspase-3 antibody staining quantification for *mrpl40* mutants at 5dpf. In all brain regions quantified, *mrpl40* mutants have fewer apoptotic cells than wildtypes.
(B) Caspase-3 antibody staining quantification for *prodha* mutants at 5dpf. In the optic tectum and forebrain, *prodha* mutants have fewer apoptotic cells than wildtypes.
Unpaired two-way Student's t-tests, p-values displayed on the plots. OT, optic tectum; FB, forebrain; HB, hindbrain.

**Supplemental Figure 3. Hindbrain proliferative zones display differences in marker gene expression**

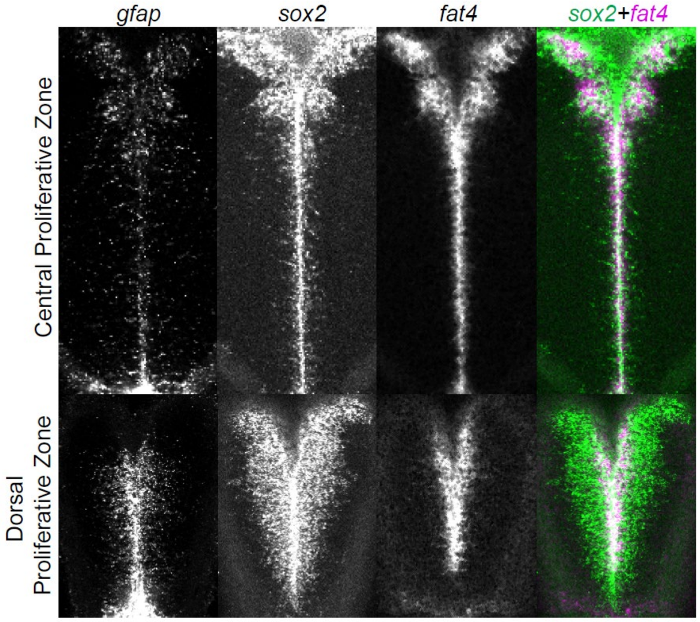


Hybridization Chain reaction *in situs* from mapzebrain (https://mapzebrain.org/home) showing that the *sox2*+ central proliferative zone is defined by *gfap/fat4* expression whereas the *sox2*+ dorsal proliferative zone lacks expression of these markers.

**Supplemental Table 2.** Morphological phenotypes observed in homozygous mutants. Related to Figure 1.

| **Mutant line** | **Details from heterozygous in-cross** | **Phenotype description of homozygotes** |
|---|---|---|
| *ess2* | Cross1: n=27/27 phenotypically normal larvae at 5dpf are het/wt. Cross2: n=48/48 phenotypically normal larvae at 5dpf are het/wt. | 3dpf: heart edema, small and curved tail; n=4/4 with phenotype are mutant |
| *ufd1l* | Cross1: n=48/48 phenotypically normal larvae at 5dpf are het/wt. Cross2: n=48/48 phenotypically normal larvae at 5dpf are het/wt. | 5dpf: curved back, dark brain; n =4/4 with phenotype are mutant |
| *cdc45* | Cross1: n=43/43 phenotypically normal larvae at 5dpf are het/wt. Cross2: n=27/41 phenotypically normal larvae at 5dpf are het/wt. | 2dpf: curved tail, small eyes, dark brain; Incompletely penetrant; n =12/12 with phenotype are mutant |
| *tbx1* | Cross1: n=41/41 phenotypically normal larvae at 5dpf are het/wt. | 3dpf: heart edema, jaw abnormalities; did not genotype phenotypic larvae as phenotype has been previously reported for *tbx1* mutants |
| *med15* | Cross1: n=35/35 phenotypically normal larvae at 5dpf are het/wt. Cross2: n=40/40 phenotypically normal larvae at 5dpf are het/wt. | 5dpf: small eyes, heart edema, jaw abnormalities; n =16/16 with phenotype are mutant |
| *C10del2* | Cross1: n=33/33 phenotypically normal larvae at 5dpf are het/wt. Cross2: n=38/38 phenotypically normal larvae at 5dpf are het/wt. | 6dpf: heart edema, small eyes, eye edema; n =25/25 with phenotype are mutant |

**Supplemental Table 3.** Incompletely penetrant swim bladder inflation phenotypes observed. Related to Figure 1.

| **Mutant line** | **Details from heterozygous in-cross** |
|---|---|
| *dgcr8* | Cross1: n=13/19 mutants without swim bladder.<br>Cross2: n=10/14 mutants without swim bladder.<br>Cross3: n=11/17 mutants without swim bladder.<br>Cross4: n=9/14 mutants without swim bladder. |
| *C5del2* | Cross1: n=7/16 mutants without swim bladder. |
| *snap29* | Cross1: n=0/7 mutants without swim bladder.<br>Cross2: n=9/11 mutants without swim bladder.<br>Cross3: n=12/12 mutants without swim bladder.<br>Cross4: n=9/11 mutants without swim bladder. |
| *C8del1* | Cross1: n=0/6 mutants without swim bladder.<br>Cross2: n=0/8 mutants without swim bladder. |
