## Supplemental Zip file 1 - python tracking scripts for "Mitochondrial genes in the 22q11.2 deleted region regulate neural stem and progenitor cell proliferation": ASR tracking file.py

```
# -*- coding: utf-8 -*-
"""
ASR Tracking code

by Philip D. Campbell


"""

import numpy as np
import cv2
#from TrackedObject import TrackedObject as TO
import matplotlib.pyplot as plt
import os
import re #regular expressions library
from skimage.measure import label, regionprops, find_contours
from skimage.morphology import remove_small_objects
from skimage import img_as_ubyte
import scipy.misc
import scipy.interpolate
import scipy.io
import pims
from tqdm import tqdm
import pandas as pd
import math
import imageio


import scipy.stats as stats
import seaborn as sns
from matplotlib.pyplot import figure
from sklearn import metrics 


#set directory containing image files
inDir = ""

# names of videos to track
Name = ''

Gene = ""

#set directory containing background files; background video used for ASR assay is itself
inDirbg = ""

# names of videos for background
Namebg = ''


#set if Tracking images to be saved
SaveTrackingIm = True; # or False

#set locations
desttracked = inDir + 'Results/TrackedVideos/' + Name + '/trackedimages/'
desttracking = inDir + 'Results/TrackedVideos/' + Name + '/'
destgene = inDir + 'Results/' + Gene + '/' + Name + '/'

# Number of fish to track
nFish = 100

#Number of rows and columns of wells
nRow = 10
nCol = 10

#Number of pixels in x and y direction
nPixel = 1024

# frames per tap event
nFrameseventtap = 100

# number of tap events
nTapevents = 100

# number of total blocks, ie how many segments to divide your data into
nBlocks = 10

#frame rate in Hz
framerate = 500

# time between frames in milliseconds
framelength = 1/framerate * 1000

# frame of tapstimulus; stimulus is after 30ms
tapstimframe = 15

# absolute total curvature in radians before considered a candidate for slc or llc, or for react
slcthreshangle = 0.8
reactthreshangle = 0.5

# time threshold for slc after stimulus, in ms; that is if movement above threshold occurs before 20ms after tap, considered an slc
slcthreshms = 20

#if no folder, make one
os.makedirs(os.path.dirname(desttracked), exist_ok=True)
os.makedirs(os.path.dirname(desttracking), exist_ok=True)
os.makedirs(os.path.dirname(destgene), exist_ok=True)


#load video
video = pims.PyAVReaderIndexed(inDir + Name)

# number of frames of video
nFrames = len(video)


#load background video
videobg = pims.PyAVReaderIndexed(inDirbg + Namebg)

# number of frames of video
nFramesbg = len(videobg)


#background subtraction
#one frame from each of the high intensity tap segments is used to create background
fr1 = videobg[2400][:,:,0]
fr2 = videobg[np.round(2900)][:,:,0]
fr3 = videobg[np.round(3400)][:,:,0]
fr4 = videobg[np.round(3900)][:,:,0]
fr5 = videobg[np.round(4400)][:,:,0]
fr6 = videobg[np.round(4900)][:,:,0]
fr7 = videobg[np.round(5400)][:,:,0]
fr8 = videobg[np.round(5900)][:,:,0]
fr9 = videobg[np.round(6400)][:,:,0]
fr10 = videobg[np.round(6900)][:,:,0]
frEnd = videobg[6999][:,:,0]

bg1 = np.median(np.stack((fr1, fr2, fr3, fr4, fr5, fr6, fr7, fr8, fr9, fr10, frEnd), axis=2), axis=2)

bg = cv2.GaussianBlur(bg1.astype(np.float64), (3,3),0)

#Define characteristics of tracking; nSeg is #points along tail, segLen1 is length in pixels to second point on tail, segLen2 is length between all other points.
#SearchAngle1 is search angle used for first segment, SearchAngle2 is search angle used for second segment, SearchAngle3 is search angle used for other segments.
#tailThresh is value above which is looked for 
nSeg = 6
segLen1 = 4.75
segLen2 = 4.75
SearchAngle1 = np.pi/2
SearchAngle2 = np.pi/2.5
SearchAngle3 = np.pi/2.25
tailThresh= 4

def tracking(bg, img, index):

    #Do a background subtraction, and blur the background
    mask = abs(bg - img)

    mask_smooth = cv2.GaussianBlur(mask, (3,3),0)

    #threshold the mask
    th_img = cv2.threshold(mask_smooth, 22, 255, cv2.THRESH_BINARY)[1]
    
    #label the connected components in the thresholded mask
    lb = label(th_img)
    
    #remove small objects from thresholded image
    lb_sorem = remove_small_objects(lb, 40)
    
    #re-label since small objects removes so get 1 to nFish
    lb2 = label(lb_sorem)

    #get properties of objects
    props = regionprops(lb2)

    # of objects    
    totobj = np.amax(lb2)
    print(str(totobj) + "in" + str(index+1))

    im_height, im_width = np.shape(mask_smooth)

    #copy the smoothed mask into a temporary image
    im_tmp = np.copy(mask_smooth)
    
    #empty arrays to hold centroid values      
    y_cent = np.zeros(nFish)
    y_cent[:] = np.nan
    x_cent = np.zeros(nFish)
    x_cent[:] = np.nan

    #empty arrays to hold points along the tail      
    pts_x = np.zeros((nFish, nSeg))
    pts_y = np.zeros((nFish, nSeg))
    pts_x[:,:] = np.nan
    pts_y[:,:] = np.nan

    #empty arrays to hold angle values     
    angles = np.zeros((nFish, nSeg))
    angles[:,:] = np.nan
    
    
    #Loop through all objects 
    for ind in range(0, totobj):

        min_row, min_col, max_row, max_col = props[ind].bbox
        
        #set curveaccum to allow to input total
        curveaccum = 0
        
        #find the centroid of fish         
        cent = props[ind].centroid
        
        #reset fishrow and fishcol tracker
        fishrow=0
        fishcol=0
        
        #go through and get row and column of this fish so can use it to index array; fish numbered from 1 in top left to  nFish in bottom right
        for row in range(1, nRow+1):

            if cent[0] < row*nPixel/nRow:
                fishrow = row
                break
        
        for col in range(1, nCol+1):
            if cent[1] < col*nPixel/nCol:
                fishcol = col
                break
        
        #this will return number of fish from 1 to nFish starting in top left corner with 1 and going L to R
        fishnum = ((fishrow-1) * nCol) + fishcol

        
        #if there is already data in arrays for this fish number, put NaN for values and go to top of for loop for next object    
        if np.isfinite(y_cent[fishnum-1]):
            y_cent[fishnum-1] = np.nan
            x_cent[fishnum-1] = np.nan
            curvature[fishnum-1,index] = np.nan
            orients[fishnum-1,index] = np.nan
            continue
        if np.isfinite(pts_y[fishnum-1,0]):
            continue
        if np.isfinite(pts_x[fishnum-1,0]):
            continue
        
        #find the centroid of the largest component
        y_cent[fishnum-1], x_cent[fishnum-1] = cent


    #get the coordinate list for the pixels in this component
        coords = props[ind].coords
    
    #Using these coordinates, search this area of the mask_smooth in order to find the 
    #point in the image with the largest diffence from the background. The coords
    #returns a list of (row,col)
    #In theory, this should be the head. Save these coordinates; np.argmax Returns 
    #the indices of the maximum values along an axis.
    
        headInd = np.argmax(mask_smooth[coords[:,0], coords[:,1]])
        y_0 = coords[headInd, 0]
        x_0 = coords[headInd, 1]


#not operator returns True if the statement is false; so if y_0 is Not A number (NaN), returns
#True and then if does not occur. if NOT NaN, returns False and if occurs
        if not np.isnan(y_0):
        
 #Assign to the first coordinates in x,y coord arrays, the centroid coordinates
            pts_x[fishnum-1,0] = x_cent[fishnum-1];
            pts_y[fishnum-1,0] = y_cent[fishnum-1];
        

        #set the head point to 0 in temporary image
            im_tmp[y_cent[fishnum-1].astype(int), x_cent[fishnum-1].astype(int)] = 0;
            

#loop over nSeg - 1 so we find (nSeg - 1) more points along the fish's back
            i = 0
            reverse = 0
            while i < nSeg-1:
            # update search direction for subsequent points
                if i > 1:
                    search_angle = np.arctan2(pts_y[fishnum-1,i]-pts_y[fishnum-1,i-1], pts_x[fishnum-1,i]-pts_x[fishnum-1,i-1])
                    ang = np.arange(search_angle-SearchAngle3,search_angle+SearchAngle3,np.pi/20)
                    angles[fishnum-1,i-1] = search_angle
                #sin(ang) * segLen will give y coordinate; cos(ang) * segLen will give x coord
                #so, this line should give several y and x values for each angle
                    ArcCoors =pts_y[fishnum-1,i] + np.sin(ang)*segLen2, pts_x[fishnum-1,i] + np.cos(ang)*segLen2
            
            
                if i == 1:
                    search_angle = np.arctan2(pts_y[fishnum-1,i]-pts_y[fishnum-1,i-1], pts_x[fishnum-1,i]-pts_x[fishnum-1,i-1])
                    ang = np.arange(search_angle-SearchAngle2,search_angle+SearchAngle2,np.pi/20)
                    angles[fishnum-1,i-1] = search_angle
                #sin(ang) * segLen will give y coordinate; cos(ang) * segLen will give x coord
                #so, this line should give several y and x values for each angle
                    ArcCoors =pts_y[fishnum-1,i] + np.sin(ang)*segLen2, pts_x[fishnum-1,i] + np.cos(ang)*segLen2
                
            # calculate search coordinates ; np.arange takes (START value, STOP value, increment)
            #Start value is included in the range, Stop value is not, will return an array
            # So, this will give an array with pi/20 spaced values btween this arc
                if i == 0:    
                    if reverse == 0:
        #define the angle, along which to search for the desired points
        #arctan2 gives the angle in radians between teh positive x-axis and the ray to point x,y
        #arctan2 changes based on sign of y, x and as such will return a value btween -pi and +pi
        # for example, if x and y<0, computes arctany/x - pi
                        search_angle = np.arctan2(y_cent[fishnum-1]-y_0, x_cent[fishnum-1] - x_0)
                    
                    if reverse == 1:
                        search_angle = np.arctan2(y_cent[fishnum-1] - pts_y[fishnum-1,1], x_cent[fishnum-1]-pts_x[fishnum-1,1])
                        
                        
                    ang = np.arange(search_angle-SearchAngle1,search_angle+SearchAngle1,np.pi/20)
                #sin(ang) * segLen will give y coordinate; cos(ang) * segLen will give x coord
                #so, this line should give several y and x values for each angle
                    ArcCoors =pts_y[fishnum-1,i] + np.sin(ang)*segLen1, pts_x[fishnum-1,i] + np.cos(ang)*segLen1
            
            #np.asarray converts input to an array with dtype whatever is specified
            # so, arccoords should be an array of coordinates to search along
                ArcCoors = np.asarray(ArcCoors, dtype=int)
            
            # avoid out of bounds issues
                ArcCoors[0][ArcCoors[0] <=0] = 1
                ArcCoors[0][ArcCoors[0] >= im_height] = im_height -1
                ArcCoors[1][ArcCoors[1] <=0] = 1
                ArcCoors[1][ArcCoors[1] >= im_width] = im_width -1

            #ArcCoords[0] refers to all y positions, [1] refers to all x positions
                if reverse == 0:    
                    im_tmp[ArcCoors[0],ArcCoors[1]] = 75
                if reverse == 1:    
                    im_tmp[ArcCoors[0],ArcCoors[1]] = 125
                    
            # find brightest pixel in smoothed image
                pt_max = np.max(mask_smooth[ArcCoors[0], ArcCoors[1]])
            
            # Returns the indices of the maximum value along an axis.
                pt_ind = np.argmax(mask_smooth[ArcCoors[0], ArcCoors[1]])
                pt_y = ArcCoors[0][pt_ind]
                pt_x = ArcCoors[1][pt_ind]


            
 # assign this as the center of the tail if the pixel is above a threshold of intesnity 
 #(this is to avoid assigning points to space not actually occupied by the fish, like if it is short, 
 #or we have gotten off track)
                if pt_max > tailThresh:
                    if reverse == 0:
                        im_tmp[pt_y.astype(int), pt_x.astype(int)] = 150 
                    if reverse == 1:
                        im_tmp[pt_y.astype(int), pt_x.astype(int)] = 250
                        
                    pts_y[fishnum-1,i+1] = pt_y
                    pts_x[fishnum-1,i+1] = pt_x

                
                    # add to curve accumlator
                    if i>1:
                        #check if angle is between 90 and 180 because run into problems when it crosses x axis
                        if abs(angles[fishnum-1,i-2]) > np.pi/2:
                            #if it is between 90 and 180, check to see if sign changes
                            if np.sign(angles[fishnum-1,i-2]) != np.sign(angles[fishnum-1,i-1]):
                                if angles[fishnum-1,i-1] > 0:
                                    curveaccum = curveaccum - ((np.pi - angles[fishnum-1,i-1]) + (np.pi + angles[fishnum-1,i-2]))
                                if angles[fishnum-1,i-1] < 0:    
                                    curveaccum = curveaccum + ((np.pi + angles[fishnum-1,i-1]) + (np.pi - angles[fishnum-1,i-2]))
                            else:
                                curveaccum = curveaccum + (angles[fishnum-1,i-1]-angles[fishnum-1,i-2])
                        else:
                            curveaccum = curveaccum + (angles[fishnum-1,i-1]-angles[fishnum-1,i-2])
                
                    if i==nSeg-2:
                       # if this is first frame of an event, should be equal to curveaccum as nothign to compare it to 
                       if index % nFrameseventtap == 0:
                            if abs(curveaccum)>np.pi/2:
                                    if reverse == 1:
                                        curvature[fishnum-1,index] = np.nan
                                        orients[fishnum-1,index] = np.nan
                                        break
                                    if reverse == 0:
                                        # switch direction of initial orientation
                                        reverse += 1
                                        # reset curveaccumulator
                                        curveaccum = 0
                                        #reset counter
                                        i = 0
                                        continue
                            else:
                                curvature[fishnum-1,index] = curveaccum
                                orients[fishnum-1,index] = angles[fishnum-1,0]
                        
                # if not first frame and if too big of a jump between frames, reverse initial direction unless this has already been done
                       else:
                           # if curveaccum is >0.8 and previous is NaN, make this one NaN as well as will not be able to tell
            #check if difference is greater than pi, if so, subtract from 2pi to get true difference
                           if np.isnan(orients[fishnum-1,index-1]):
                               curvature[fishnum-1,index] = np.nan
                               orients[fishnum-1,index] = np.nan
                               break
                           
                           orients[fishnum-1,index] = angles[fishnum-1,0]
                            
                           if abs(orients[fishnum-1,index-1] - orients[fishnum-1,index]) > np.pi:
                                deltaorient = (2*np.pi) - abs(orients[fishnum-1,index-1] - orients[fishnum-1,index])
                           else:
                                deltaorient = abs(orients[fishnum-1,index-1] - orients[fishnum-1,index])
                           
                           if deltaorient > np.pi/2:  
                                    print(str(fishnum) + " for " + str(deltaorient) + " in index " + str(index))
                                    if reverse == 1:
                                        curvature[fishnum-1,index] = np.nan
                                        #print(str(fishnum) + " for " + str(curvature[fishnum-1]) + " in i " + str(i))
                                        orients[fishnum-1,index] = np.nan
                                        break
                                    if reverse == 0:
                                        # switch direction of initial orientation
                                        reverse += 1
                                        # reset curveaccumulator
                                        curveaccum = 0
                                        #reset counter
                                        i = 0
                                        continue
                           else:
                                    curvature[fishnum-1,index] = curveaccum
                                    orients[fishnum-1,index] = angles[fishnum-1,0]

                    i += 1
                
                else: #  if we didnt find enough point on the tail, try to reverse initial search unless this has already been done
                    if i <= (nSeg-2):
                                        
                        if index % nFrameseventtap == 0:
                        
                            if reverse == 1:
                                curvature[fishnum-1,index] = np.nan
                                orients[fishnum-1, index] = np.nan
                                break
                            if reverse == 0:
                                # switch direction of initial orientation
                                reverse += 1
                                # reset curveaccumulator
                                curveaccum = 0
                                #reset counter
                                i = 0
                                continue
                        else:
                            if np.isnan(orients[fishnum-1,index-1]):
                               curvature[fishnum-1,index] = np.nan
                               orients[fishnum-1,index] = np.nan
                               break                            
                            
                            orients[fishnum-1,index] = angles[fishnum-1,0]
                    
                            if abs(orients[fishnum-1,index-1] - orients[fishnum-1,index]) > np.pi:
                                deltaorient = (2*np.pi) - abs(orients[fishnum-1,index-1] - orients[fishnum-1,index])
                            else:
                                deltaorient = abs(orients[fishnum-1,index-1] - orients[fishnum-1,index])
                           
                            if deltaorient > np.pi/2:  
                                if reverse == 1:
                                    curvature[fishnum-1,index] = np.nan
                                    orients[fishnum-1,index] = np.nan
                                    break
                                if reverse == 0:
                                    # switch direction of initial orientation
                                    reverse += 1
                                    # reset curveaccumulator
                                    curveaccum = 0
                                    #reset counter
                                    i = 0
                                    continue
                            else:
                                curvature[fishnum-1,index] = np.nan
                                orients[fishnum-1, index] = angles[fishnum-1,0]
                                break
                
                    else:
                        curvature[fishnum-1,index] = np.nan
                        orients[fishnum-1, index] = angles[fishnum-1,0]
                        break
                    
    return im_tmp, pts_y, pts_x, y_cent, x_cent, curvature, orients


#Initialize arrays to hold values
xycoords = np.zeros((nFish, nSeg, 2, nFrames))
cents = np.zeros((nFish, 2, nFrames))
curvature = np.zeros((nFish, nFrames))
orients = np.zeros((nFish, nFrames))
orients[:] = np.nan

curvature[:] = np.nan

#tracking

print("Tracking....")
    
#tqdm shows a progress bar around any iterable
for index, img in enumerate(tqdm(video)):
       
        if (index < nFrames):

            img = img[:,:,0] # its actually a black and white image, but gets read in as three channel
            
            # blur image
            img = cv2.GaussianBlur(img.astype(np.float64), (3,3),0)
            
            #calling the tracking image with bg and individual img
            timg, pts_y, pts_x, y_cent, x_cent, curvature, orients = tracking(bg, img, index)
            
            cents[:, 0, index] = y_cent
            cents[:, 1, index] = x_cent
            xycoords[:, :, 0, index] = pts_y
            xycoords[:, :, 1, index] = pts_x

            #save the tracked coordinates and the tracked image files

            if SaveTrackingIm:

                cv2.imwrite(inDir + 'Results/TrackedVideos/' + Name + '/trackedimages/' + Name + '_' + str(index) +'.png', np.vstack((timg)))         
     
        
print("Saving tracking data arrays....")
np.savez(inDir + 'Results/TrackedVideos/' + Name + '/TrackData.npz', xycoords=xycoords, cents=cents, curvature=curvature, orients=orients)
        
import seaborn as sns
from scipy.signal import medfilt

# START CALCULATIONS HERE

#setup med-filtered arrays
medfiltcentY = np.zeros((nFish, nFrames))
medfiltcentY[:,:] = np.nan
medfiltcentX = np.zeros((nFish, nFrames))
medfiltcentX[:,:] = np.nan


for number in range(nFish):
    for i in range(0,nTapevents):
        medfiltcentY[number,(i*nFrameseventtap):((i+1)*nFrameseventtap)] = medfilt(cents[number,0,(i*nFrameseventtap):((i+1)*nFrameseventtap)],3)
        medfiltcentX[number,(i*nFrameseventtap):((i+1)*nFrameseventtap)] = medfilt(cents[number,1,(i*nFrameseventtap):((i+1)*nFrameseventtap)],3)
    
    
    
# number of event blocks for sens
nBlockssens = 2

# number of ppi types
nBlocksppi = 5

# number of event blocks for sens
nBlockshab = 3


#set up arrays for data
disp = np.zeros((nFish, nFrames-1))
disp[:,:] = np.nan
filtdisp = np.zeros((nFish, nFrames-1))
filtdisp[:,:] = np.nan

angdisp = np.zeros((nFish, nFrames-1))
angdisp[:,:] = np.nan

angdispmax = np.zeros((nFish, nTapevents))
angdispmax[:,:] = np.nan


latencytaps = np.zeros((nFish, nTapevents))
latencytaps[:,:] = np.nan
latencytapsms = np.zeros((nFish, nTapevents))
latencytapsms[:,:] = np.nan 
slctaps = np.zeros((nFish, nTapevents))
slctaps[:,:] = np.nan
llctaps = np.zeros((nFish, nTapevents))
llctaps[:,:] = np.nan
reacttaps = np.zeros((nFish, nTapevents))
reacttaps[:,:] = np.nan
nomvmttaps = np.zeros((nFish, nTapevents))
nomvmttaps[:,:] = np.nan
tapbendmax = np.zeros((nFish, nTapevents))
tapbendmax[:,:] = np.nan
tapdeltaorient = np.zeros((nFish, nTapevents))
tapdeltaorient[:,:] = np.nan
tapdisplacement = np.zeros((nFish, nTapevents))
tapdisplacement[:,:] = np.nan
tapdistance = np.zeros((nFish, nTapevents))
tapdistance[:,:] = np.nan
movementyesno = np.zeros((nFish, nFrames-1))
movementyesno[:,:] = np.nan
movementyesnofill = np.zeros((nFish, nFrames-1))
movementyesnofill[:,:] = np.nan
tapduration = np.zeros((nFish, nTapevents))
tapduration[:,:] = np.nan

tapright = np.zeros((nFish, nTapevents))
tapright[:,:] = np.nan
taptowards = np.zeros((nFish, nTapevents))
taptowards[:,:] = np.nan
taptowardshalf = np.zeros((nFish, nTapevents))
taptowardshalf[:,:] = np.nan

print("Calculating displacement....")
# calculating displacement and thigmotaxis
for number in range(nFish):
    i = 0
    while i < nFrames:
        if  np.isnan(cents[number, 0, i:((math.floor(i/nFrameseventtap)+1)*nFrameseventtap)]).any():
            #if no centroid is defined in any frame of this event, skip to next block
            event = math.floor(i/nFrameseventtap)
            i = (event+1)*nFrameseventtap
            continue

        else:            

            #calculate displacement between each frame
            if i>0:
                disp[number,i-1] = np.sqrt((cents[number, 0, i]-cents[number, 0, i-1])**2 + (cents[number, 1, i]-cents[number, 1, i-1])**2) 
                filtdisp[number,i-1] = np.sqrt((medfiltcentY[number, i]-medfiltcentY[number, i-1])**2 + (medfiltcentX[number, i]-medfiltcentX[number, i-1])**2) 
                   # define movement as displacement >0.2 and populate array with 1 for >0.2 and 0 for <0.2; nan if displ nan
                if filtdisp[number,i-1] > 0.5:
                    movementyesno[number,i-1] = 1
                else:
                    movementyesno[number,i-1] = 0
            
            i=i+1

print("Calculating angular displacement....")
# calculating displacement and thigmotaxis
for number in range(nFish):
    i = 0
    while i < nFrames:
        if  np.isnan(orients[number, i:((math.floor(i/nFrameseventtap)+1)*nFrameseventtap)]).any():
            #if no orient is defined in any frame of this event, skip to next block
            event = math.floor(i/nFrameseventtap)
            i = (event+1)*nFrameseventtap
            continue

        else:            
            #calculate angular displacement between each frame
            if i>0:

            #check if difference is greater than pi, if so, subtract from 2pi to get true difference
                if abs(orients[number, i] - orients[number, i-1]) > np.pi:
                    angdisp[number,i-1] = (2*np.pi) - abs(orients[number, i] - orients[number, i-1])
                else:
                    angdisp[number,i-1] = abs(orients[number, i] - orients[number, i-1])
            i=i+1

print("Defining movement....")
#Attempt to fill holes of movement array. Will eliminate singleton 1s or 0s so that for movement will need >1 frame with value of
                       #1 and to stop movement will need >1 frame with value of 0
# fill initial frame with values of initial array
movementyesnofill[:, 0] = movementyesno[:, 0]

for number in range(nFish):
    for i in range(1,nFrames-1):
        if  np.isnan(movementyesno[number, i]):
            continue
        #if nan either before or after value i, just keep value because cannot tell
        if  np.isnan(movementyesnofill[number, i-1]):
            movementyesnofill[number, i] = movementyesno[number, i]
            continue
        if i < nFrames-2:
            if np.isnan(movementyesno[number, i+1]):
                movementyesnofill[number, i] = movementyesno[number, i]
                continue


# here is where we define movement and stopping as >1 frame of either 1 or 0
        if movementyesno[number, i] == 0:
            if movementyesnofill[number, i-1] == 0:
                movementyesnofill[number, i] = 0
            if movementyesnofill[number, i-1] == 1:
                if i < nFrames-2:
                    if movementyesno[number, i+1] == 1:
                        movementyesnofill[number, i] = 1
                    if movementyesno[number, i+1] == 0:
                        movementyesnofill[number, i] = 0
                else:
                    movementyesnofill[number, i] = 0
        
        if movementyesno[number, i] == 1:
            if movementyesnofill[number, i-1] == 1:
                movementyesnofill[number, i] = 1
            if movementyesnofill[number, i-1] == 0:
                if i < nFrames-2:
                    if movementyesno[number, i+1] == 0:
                        movementyesnofill[number, i] = 0
                    if movementyesno[number, i+1] == 1:
                        movementyesnofill[number, i] = 1
                else:
                    movementyesnofill[number, i] = 1
                    
                    


print("Finding tap latency and Classifying SLC, LLC....")
#finding tap response frame and latency
for number in range(nFish):
    i=0
    while i < nFrames:
        # calculating latency to bend
        
        # if this frame and next frame have curvatures of NaN, eliminate this event - make latency Nan, then skip to next event
        if np.isnan(curvature[number, i]) and np.isnan(curvature[number, i+1]) :
            event = math.floor(i/nFrameseventtap)
            latencytaps[number,event] = np.nan
            i = (event+1)*nFrameseventtap
            continue
        # if only this frame has curvature of Nan, go to next frame
        if np.isnan(curvature[number, i]):
            i = i+1
            continue
        else:    
            event = math.floor(i/nFrameseventtap)

            if np.nanmax(abs(curvature[number, ((event*nFrameseventtap) + tapstimframe):((event+1)*nFrameseventtap)])) > slcthreshangle:
                if abs(curvature[number, i]) > slcthreshangle:
                    latencytaps[number,event] = i - (nFrameseventtap*event)
                    if latencytaps[number,event] <= tapstimframe:
                        latencytaps[number,event] = np.nan
                    if latencytaps[number,event] > tapstimframe and latencytaps[number,event]<=(tapstimframe+ (slcthreshms/framelength)):
                        slctaps[number,event] = 1
                    if latencytaps[number,event] > (tapstimframe + (slcthreshms/framelength)):
                        llctaps[number,event] = 1
                    i = (event+1)*nFrameseventtap
                    continue
                else:
                    if (i+1) % nFrameseventtap == 0:
                        nomvmttaps[number,event] = 1
                        latencytaps[number,event] = np.nan
                    i = i+1
                    continue
            if np.nanmax(abs(curvature[number, ((event*nFrameseventtap) + tapstimframe):((event+1)*nFrameseventtap)])) > reactthreshangle:       
                if abs(curvature[number, i]) > reactthreshangle:
                    latencytaps[number,event] = i - (nFrameseventtap*event)
                    if latencytaps[number,event] <= tapstimframe:
                        latencytaps[number,event] = np.nan
                    else:
                        #classify as react
                        reacttaps[number,event] = 1
                    i = (event+1)*nFrameseventtap
                    continue
                else:
                    if (i+1) % nFrameseventtap == 0:
                        nomvmttaps[number,event] = 1
                        latencytaps[number,event] = np.nan
                    i = i+1
                    continue
            else:
                #classify as no movement
                nomvmttaps[number,event] = 1
                latencytaps[number,event] = np.nan
                i = (event+1)*nFrameseventtap

# calculate latency in ms from frame number where > certain radian
latencytapsms[:,:] = (latencytaps[:,:]*framelength) - (tapstimframe*framelength)

print("Calculating bend max, delta orient, total displacement....")                     
# calculating bend max, change in orientation, and totaldisplacement for taps
for number in range(nFish):
    i=0
    while i < nTapevents:
        #check if latency is before stimulus (ie. nan)
        if np.isnan(latencytaps[number,i]):
            i=i+1
            continue
        
        else:
            #time moving
            tapduration[number,i] = np.sum(movementyesnofill[number,((i*nFrameseventtap)+tapstimframe):(((i+1)*nFrameseventtap)-1)])*framelength
            
            #label bouts in block
            lb = label(movementyesnofill[number, ((i*nFrameseventtap)+tapstimframe):(((i+1)*nFrameseventtap)-1)])
            
            #remove bouts less than 2 - though shouldn't be any anyways
            lb_sorem = remove_small_objects(lb, 2)
            
            #re-label since small objects removed
            lb2 = label(lb_sorem)
                      
            #define array of distances in this block
            x = filtdisp[number, ((i*nFrameseventtap)+tapstimframe):(((i+1)*nFrameseventtap)-1)]
            
            #total distance: is equal to sum of displacement following tap stimulus where movementyesnofill is 1
            tapdistance[number,i] = np.sum(x[lb2!=0])

           
            #change in orientation
            #calculate frame right before tap
            tapframe = i*nFrameseventtap + (tapstimframe-1)
            
            #check if difference is greater than pi, if so, subtract from 2pi to get true difference
            if abs(orients[number, (nFrameseventtap*(i+1))-1] - orients[number, tapframe]) > np.pi:
                tapdeltaorient[number, i] = (2*np.pi) - abs(orients[number, (nFrameseventtap*(i+1))-1] - orients[number, tapframe])
            else:
                tapdeltaorient[number, i] = abs(orients[number, (nFrameseventtap*(i+1))-1] - orients[number, tapframe])
            
            #tap displacement
            tapdisplacement[number,i] = np.sqrt((cents[number, 0, tapframe]-cents[number, 0, ((i+1)*nFrameseventtap)-1])**2 + (cents[number, 1, tapframe]-cents[number, 1, ((i+1)*nFrameseventtap)-1])**2)
            
            
            # for react or nomovement cases, just check for max bend angle after tap
            if latencytaps[number,i] == (nFrameseventtap-1):
                tapbendmax[number,i] = np.nanmax(abs(curvature[number, ((i*nFrameseventtap) + tapstimframe):((i+1)*nFrameseventtap)]))
                i=i+1
            else:

                #for other bend max: calculate frame where above threshold movement happens
                frameorient = i*nFrameseventtap + (latencytaps[number,i]) - 5
                framecurve = i*nFrameseventtap + (latencytaps[number,i])
                
            #compute L/R and towards/away; if orientation of fish is between 0 and pi
                if orients[number, int(frameorient)] > 0:
                    #if curvature is negative, meaning CW
                    if curvature[number,int(framecurve)] < 0:
                        tapright[number,i] = 1
                        taptowards[number,i] = 1
                        
                    #if curvature is positive, meaning CCW
                    if curvature[number,int(framecurve)] > 0:
                        tapright[number,i] = 0
                        taptowards[number,i] = 0
            #if initial orient is within middle 90deg
                    if abs(orients[number, int(frameorient)]) > (4.5*np.pi/18) and abs(orients[number, int(frameorient)]) < (13.5*np.pi/18):
                        if curvature[number,int(framecurve)] < 0:
                            taptowardshalf[number,i] = 1
                        
                        if curvature[number,int(framecurve)] > 0:
                            taptowardshalf[number,i] = 0    

            
            #if orientation of fish is between 0 and pi
                if orients[number, int(frameorient)] < 0:
                    #if curvature is negative, meaning CW
                    if curvature[number,int(framecurve)] < 0:
                        tapright[number,i] = 1
                        taptowards[number,i] = 0
                        
                    #if curvature is positive, meaning CCW
                    if curvature[number,int(framecurve)] > 0:
                        tapright[number,i] = 0
                        taptowards[number,i] = 1

            #if initial orient is within middle 90deg
                    if abs(orients[number, int(frameorient)]) > (4.5*np.pi/18) and abs(orients[number, int(frameorient)]) < (13.5*np.pi/18):
                        if curvature[number,int(framecurve)] < 0:
                            taptowardshalf[number,i] = 0
                        
                        if curvature[number,int(framecurve)] > 0:
                            taptowardshalf[number,i] = 1  

                #for other bend max: calculate frame where above threshold movement happens
                frame = i*nFrameseventtap + (latencytaps[number,i])
                j=i
                if frame+2>nFrames-1:
                    i=i+1
                    continue
                else:                 
                
                    while frame < (j+1)*nFrameseventtap:
                #check if curvature is nan around bend max, if so break while loop as will not be able to decipher bendmax
                        if np.isnan(curvature[number,int(frame)]) or np.isnan(curvature[number,int(frame-1)]) or np.isnan(curvature[number,int(frame+1)]) or np.isnan(curvature[number,int(frame+2)]):
                            j=j+1
                            break
                        else:
                    # find point of local max right after threshold reached
                            if abs(curvature[number,int(frame)]) > abs(curvature[number,int(frame-1)]) and abs(curvature[number,int(frame)]) > abs(curvature[number,int(frame+1)]) and abs(curvature[number,int(frame)]) > abs(curvature[number,int(frame+2)]):
                                tapbendmax[number,j] = curvature[number,int(frame)]
                                j=j+1
                                break
                    # if at end of tap window, move on to next event
                            else:
                                frame = frame + 1
                                if frame == ((j+1)*nFrameseventtap)-2:
                                    j=j+1
                                    break
                
                frame = i*nFrameseventtap + (latencytaps[number,i])
                if frame+3>nFrames-1:
                    i=i+1
                    continue
                else:
                
                    while frame < (i+1)*nFrameseventtap:
                        #check if curvature is nan around bend max, if so break while loop as will not be able to decipher bendmax
                        if np.isnan(angdisp[number,int(frame)]) or np.isnan(angdisp[number,int(frame-1)]) or np.isnan(angdisp[number,int(frame-2)]) or np.isnan(angdisp[number,int(frame-3)]):
                            i=i+1
                            break
                        else:
                    # find point of local max right after threshold reached
                            if abs(angdisp[number,int(frame-2)]) > abs(angdisp[number,int(frame-3)]) and abs(angdisp[number,int(frame-2)]) > abs(angdisp[number,int(frame-1)]) and abs(angdisp[number,int(frame-2)]) > abs(angdisp[number,int(frame)]):
                                angdispmax[number,i] = angdisp[number,int(frame-2)]
                                i=i+1
                                break
                    # if at end of tap window, move on to next event
                            else:
                                frame = frame + 1
                                if frame == ((i+1)*nFrameseventtap):
                                    i=i+1
                                    break
 
print("Setting up final arrays....")   

slcnan = np.zeros((nFish, nBlocks))
slcnan[:] = np.nan
slctot = np.zeros((nFish, nBlocks))
slctot[:] = np.nan
latnan = np.zeros((nFish, nBlocks))
latnan[:] = np.nan
lattot = np.zeros((nFish, nBlocks))
lattot[:] = np.nan
tapbendmaxnan = np.zeros((nFish, nBlocks))
tapbendmaxnan[:] = np.nan
tapbendmaxtot = np.zeros((nFish, nBlocks))
tapbendmaxtot[:] = np.nan
tapdeltaorientnan = np.zeros((nFish, nBlocks))
tapdeltaorientnan[:] = np.nan
tapdeltaorienttot = np.zeros((nFish, nBlocks))
tapdeltaorienttot[:] = np.nan
tapdisplacementnan = np.zeros((nFish, nBlocks))
tapdisplacementnan[:] = np.nan
tapdisplacementtot = np.zeros((nFish, nBlocks))
tapdisplacementtot[:] = np.nan
tapdistancenan = np.zeros((nFish, nBlocks))
tapdistancenan[:] = np.nan
tapdistancetot = np.zeros((nFish, nBlocks))
tapdistancetot[:] = np.nan
angdispmaxnan = np.zeros((nFish, nBlocks))
angdispmaxnan[:] = np.nan
angdispmaxtot = np.zeros((nFish, nBlocks))
angdispmaxtot[:] = np.nan


avglat = np.zeros((nFish, nBlocks))
avglat[:] = np.nan
medlat = np.zeros((nFish, nBlocks))
medlat[:] = np.nan
slc = np.zeros((nFish, nBlocks))
slc[:] = np.nan
llc = np.zeros((nFish, nBlocks))
llc[:] = np.nan
react = np.zeros((nFish, nBlocks))
react[:] = np.nan
nomvmt = np.zeros((nFish, nBlocks))
nomvmt[:] = np.nan
avgbend = np.zeros((nFish, nBlocks))
avgbend[:] = np.nan
medbend = np.zeros((nFish, nBlocks))
medbend[:] = np.nan
avgorient = np.zeros((nFish, nBlocks))
avgorient[:] = np.nan
medorient = np.zeros((nFish, nBlocks))
medorient[:] = np.nan
avgdisp = np.zeros((nFish, nBlocks))
avgdisp[:] = np.nan
meddisp = np.zeros((nFish, nBlocks))
meddisp[:] = np.nan
avgdist = np.zeros((nFish, nBlocks))
avgdist[:] = np.nan
meddist = np.zeros((nFish, nBlocks))
meddist[:] = np.nan
meddist[:] = np.nan
avgduration = np.zeros((nFish, nBlocks))
avgduration[:] = np.nan
medduration = np.zeros((nFish, nBlocks))
medduration[:] = np.nan
avgangvelmax = np.zeros((nFish, nBlocks))
avgangvelmax[:] = np.nan
medangvelmax = np.zeros((nFish, nBlocks))
medangvelmax[:] = np.nan

right = np.zeros((nFish, nBlocks))
right[:] = np.nan
towards = np.zeros((nFish, nBlocks))
towards[:] = np.nan
towardshalf = np.zeros((nFish, nBlocks))
towardshalf[:] = np.nan


print("Tabulating results....")  
#count number of nan and total events and make calculations for sensitivity events
for number in range(nFish):
    for i in range(nBlockssens):
        slcnan[number,i] = np.count_nonzero(np.isnan(latencytaps[number, int(i*(nTapevents/nBlocks)):int((i+1)*(nTapevents/nBlocks))])) - np.count_nonzero(np.isfinite(nomvmttaps[number, int(i*(nTapevents/nBlocks)):int((i+1)*(nTapevents/nBlocks))]))
        slctot[number,i] = np.size(latencytaps[number, int(i*(nTapevents/nBlocks)):int((i+1)*(nTapevents/nBlocks))])
        latnan[number,i] = np.count_nonzero(np.isnan(latencytaps[number, int(i*(nTapevents/nBlocks)):int((i+1)*(nTapevents/nBlocks))]))
        lattot[number,i] = np.size(latencytaps[number, int(i*(nTapevents/nBlocks)):int((i+1)*(nTapevents/nBlocks))])
        tapbendmaxnan[number,i] = np.count_nonzero(np.isnan(tapbendmax[number, int(i*(nTapevents/nBlocks)):int((i+1)*(nTapevents/nBlocks))]))
        tapbendmaxtot[number,i] = np.size(tapbendmax[number, int(i*(nTapevents/nBlocks)):int((i+1)*(nTapevents/nBlocks))])
        tapdeltaorientnan[number,i] = np.count_nonzero(np.isnan(tapdeltaorient[number, int(i*(nTapevents/nBlocks)):int((i+1)*(nTapevents/nBlocks))]))
        tapdeltaorienttot[number,i] = np.size(tapdeltaorient[number, int(i*(nTapevents/nBlocks)):int((i+1)*(nTapevents/nBlocks))])
        tapdisplacementnan[number,i] = np.count_nonzero(np.isnan(tapdisplacement[number, int(i*(nTapevents/nBlocks)):int((i+1)*(nTapevents/nBlocks))]))
        tapdisplacementtot[number,i] = np.size(tapdisplacement[number, int(i*(nTapevents/nBlocks)):int((i+1)*(nTapevents/nBlocks))])
        tapdistancenan[number,i] = np.count_nonzero(np.isnan(tapdistance[number, int(i*(nTapevents/nBlocks)):int((i+1)*(nTapevents/nBlocks))]))
        tapdistancetot[number,i] = np.size(tapdistance[number, int(i*(nTapevents/nBlocks)):int((i+1)*(nTapevents/nBlocks))])
        angdispmaxnan[number,i] = np.count_nonzero(np.isnan(angdispmax[number, int(i*(nTapevents/nBlocks)):int((i+1)*(nTapevents/nBlocks))]))
        angdispmaxtot[number,i] = np.size(angdispmax[number, int(i*(nTapevents/nBlocks)):int((i+1)*(nTapevents/nBlocks))])

        avglat[number,i] = np.nanmean(latencytapsms[number, int(i*(nTapevents/nBlocks)):int((i+1)*(nTapevents/nBlocks))])
        medlat[number,i] = np.nanmedian(latencytapsms[number, int(i*(nTapevents/nBlocks)):int((i+1)*(nTapevents/nBlocks))])
        slc[number,i] = np.nansum(slctaps[number, int(i*(nTapevents/nBlocks)):int((i+1)*(nTapevents/nBlocks))])/(slctot[number,i]-slcnan[number,i])
        llc[number,i] = np.nansum(llctaps[number, int(i*(nTapevents/nBlocks)):int((i+1)*(nTapevents/nBlocks))])/(slctot[number,i]-slcnan[number,i])
        react[number,i] = np.nansum(reacttaps[number, int(i*(nTapevents/nBlocks)):int((i+1)*(nTapevents/nBlocks))])/(slctot[number,i]-slcnan[number,i])
        nomvmt[number,i] = np.nansum(nomvmttaps[number, int(i*(nTapevents/nBlocks)):int((i+1)*(nTapevents/nBlocks))])/(slctot[number,i]-slcnan[number,i])
        
        right[number,i] = np.nansum(tapright[number, int(i*(nTapevents/nBlocks)):int((i+1)*(nTapevents/nBlocks))])/(lattot[number,i]-latnan[number,i])
        towards[number,i] = np.nansum(taptowards[number, int(i*(nTapevents/nBlocks)):int((i+1)*(nTapevents/nBlocks))])/(lattot[number,i]-latnan[number,i])
        towardshalf[number,i] = np.nansum(taptowardshalf[number, int(i*(nTapevents/nBlocks)):int((i+1)*(nTapevents/nBlocks))])/np.count_nonzero(np.isfinite(taptowardshalf[number, int(i*(nTapevents/nBlocks)):int((i+1)*(nTapevents/nBlocks))]))

        
        avgbend[number,i] = np.nanmean(abs(tapbendmax[number, int(i*(nTapevents/nBlocks)):int((i+1)*(nTapevents/nBlocks))]))
        medbend[number,i] = np.nanmedian(abs(tapbendmax[number, int(i*(nTapevents/nBlocks)):int((i+1)*(nTapevents/nBlocks))]))
        avgorient[number,i] = np.nanmean(abs(tapdeltaorient[number, int(i*(nTapevents/nBlocks)):int((i+1)*(nTapevents/nBlocks))]))
        medorient[number,i] = np.nanmedian(abs(tapdeltaorient[number, int(i*(nTapevents/nBlocks)):int((i+1)*(nTapevents/nBlocks))]))
        avgdisp[number,i] = np.nanmean(abs(tapdisplacement[number, int(i*(nTapevents/nBlocks)):int((i+1)*(nTapevents/nBlocks))]))
        meddisp[number,i] = np.nanmedian(abs(tapdisplacement[number, int(i*(nTapevents/nBlocks)):int((i+1)*(nTapevents/nBlocks))]))
        avgdist[number,i] = np.nanmean(abs(tapdistance[number, int(i*(nTapevents/nBlocks)):int((i+1)*(nTapevents/nBlocks))]))
        meddist[number,i] = np.nanmedian(abs(tapdistance[number, int(i*(nTapevents/nBlocks)):int((i+1)*(nTapevents/nBlocks))]))
        avgduration[number,i] = np.nanmean(abs(tapduration[number, int(i*(nTapevents/nBlocks)):int((i+1)*(nTapevents/nBlocks))]))
        medduration[number,i] = np.nanmedian(abs(tapduration[number, int(i*(nTapevents/nBlocks)):int((i+1)*(nTapevents/nBlocks))]))
        avgangvelmax[number,i] = np.nanmean(abs(angdispmax[number, int(i*(nTapevents/nBlocks)):int((i+1)*(nTapevents/nBlocks))]))
        medangvelmax[number,i] = np.nanmedian(abs(angdispmax[number, int(i*(nTapevents/nBlocks)):int((i+1)*(nTapevents/nBlocks))]))        


#count number of nan and total events and make calculations for ppi events
for number in range(nFish):
    for i in range(nBlocksppi):
        slcnan[number,(nBlockssens+i)] = np.count_nonzero(np.isnan(latencytaps[number, [*range(int(((nBlockssens)*(nTapevents/nBlocks))+i),int((nBlockssens+nBlocksppi)*(nTapevents/nBlocks)),nBlocksppi)]])) - np.count_nonzero(np.isfinite(nomvmttaps[number, [*range(int(((nBlockssens)*(nTapevents/nBlocks))+i),int((nBlockssens+nBlocksppi)*(nTapevents/nBlocks)),nBlocksppi)]]))
        slctot[number,(nBlockssens+i)] = np.size(latencytaps[number, [*range(int(((nBlockssens)*(nTapevents/nBlocks))+i),int((nBlockssens+nBlocksppi)*(nTapevents/nBlocks)),nBlocksppi)]])
        latnan[number,(nBlockssens+i)] = np.count_nonzero(np.isnan(latencytaps[number, [*range(int(((nBlockssens)*(nTapevents/nBlocks))+i),int((nBlockssens+nBlocksppi)*(nTapevents/nBlocks)),nBlocksppi)]]))
        lattot[number,(nBlockssens+i)] = np.size(latencytaps[number, [*range(int(((nBlockssens)*(nTapevents/nBlocks))+i),int((nBlockssens+nBlocksppi)*(nTapevents/nBlocks)),nBlocksppi)]])
        tapbendmaxnan[number,(nBlockssens+i)] = np.count_nonzero(np.isnan(tapbendmax[number, [*range(int(((nBlockssens)*(nTapevents/nBlocks))+i),int((nBlockssens+nBlocksppi)*(nTapevents/nBlocks)),nBlocksppi)]]))
        tapbendmaxtot[number,(nBlockssens+i)] = np.size(tapbendmax[number, [*range(int(((nBlockssens)*(nTapevents/nBlocks))+i),int((nBlockssens+nBlocksppi)*(nTapevents/nBlocks)),nBlocksppi)]])
        tapdeltaorientnan[number,(nBlockssens+i)] = np.count_nonzero(np.isnan(tapdeltaorient[number, [*range(int(((nBlockssens)*(nTapevents/nBlocks))+i),int((nBlockssens+nBlocksppi)*(nTapevents/nBlocks)),nBlocksppi)]]))
        tapdeltaorienttot[number,(nBlockssens+i)] = np.size(tapdeltaorient[number, [*range(int(((nBlockssens)*(nTapevents/nBlocks))+i),int((nBlockssens+nBlocksppi)*(nTapevents/nBlocks)),nBlocksppi)]])
        tapdisplacementnan[number,(nBlockssens+i)] = np.count_nonzero(np.isnan(tapdisplacement[number, [*range(int(((nBlockssens)*(nTapevents/nBlocks))+i),int((nBlockssens+nBlocksppi)*(nTapevents/nBlocks)),nBlocksppi)]]))
        tapdisplacementtot[number,(nBlockssens+i)] = np.size(tapdisplacement[number, [*range(int(((nBlockssens)*(nTapevents/nBlocks))+i),int((nBlockssens+nBlocksppi)*(nTapevents/nBlocks)),nBlocksppi)]])
        tapdistancenan[number,(nBlockssens+i)] = np.count_nonzero(np.isnan(tapdistance[number, [*range(int(((nBlockssens)*(nTapevents/nBlocks))+i),int((nBlockssens+nBlocksppi)*(nTapevents/nBlocks)),nBlocksppi)]]))
        tapdistancetot[number,(nBlockssens+i)] = np.size(tapdistance[number, [*range(int(((nBlockssens)*(nTapevents/nBlocks))+i),int((nBlockssens+nBlocksppi)*(nTapevents/nBlocks)),nBlocksppi)]])
        angdispmaxnan[number,(nBlockssens+i)] = np.count_nonzero(np.isnan(angdispmax[number, [*range(int(((nBlockssens)*(nTapevents/nBlocks))+i),int((nBlockssens+nBlocksppi)*(nTapevents/nBlocks)),nBlocksppi)]]))
        angdispmaxtot[number,(nBlockssens+i)] = np.size(angdispmax[number, [*range(int(((nBlockssens)*(nTapevents/nBlocks))+i),int((nBlockssens+nBlocksppi)*(nTapevents/nBlocks)),nBlocksppi)]])


        avglat[number,(nBlockssens+i)] = np.nanmean(latencytapsms[number, [*range(int(((nBlockssens)*(nTapevents/nBlocks))+i),int((nBlockssens+nBlocksppi)*(nTapevents/nBlocks)),nBlocksppi)]])
        medlat[number,(nBlockssens+i)] = np.nanmedian(latencytapsms[number, [*range(int(((nBlockssens)*(nTapevents/nBlocks))+i),int((nBlockssens+nBlocksppi)*(nTapevents/nBlocks)),nBlocksppi)]])
        slc[number,(nBlockssens+i)] = np.nansum(slctaps[number, [*range(int(((nBlockssens)*(nTapevents/nBlocks))+i),int((nBlockssens+nBlocksppi)*(nTapevents/nBlocks)),nBlocksppi)]])/(slctot[number,(nBlockssens+i)]-slcnan[number,(nBlockssens+i)])
        llc[number,(nBlockssens+i)] = np.nansum(llctaps[number, [*range(int(((nBlockssens)*(nTapevents/nBlocks))+i),int((nBlockssens+nBlocksppi)*(nTapevents/nBlocks)),nBlocksppi)]])/(slctot[number,(nBlockssens+i)]-slcnan[number,(nBlockssens+i)])
        react[number,(nBlockssens+i)] = np.nansum(reacttaps[number, [*range(int(((nBlockssens)*(nTapevents/nBlocks))+i),int((nBlockssens+nBlocksppi)*(nTapevents/nBlocks)),nBlocksppi)]])/(slctot[number,(nBlockssens+i)]-slcnan[number,(nBlockssens+i)])
        nomvmt[number,(nBlockssens+i)] = np.nansum(nomvmttaps[number, [*range(int(((nBlockssens)*(nTapevents/nBlocks))+i),int((nBlockssens+nBlocksppi)*(nTapevents/nBlocks)),nBlocksppi)]])/(slctot[number,(nBlockssens+i)]-slcnan[number,(nBlockssens+i)])

        right[number,(nBlockssens+i)] = np.nansum(tapright[number, [*range(int(((nBlockssens)*(nTapevents/nBlocks))+i),int((nBlockssens+nBlocksppi)*(nTapevents/nBlocks)),nBlocksppi)]])/(lattot[number,(nBlockssens+i)]-latnan[number,(nBlockssens+i)])
        towards[number,(nBlockssens+i)] = np.nansum(taptowards[number, [*range(int(((nBlockssens)*(nTapevents/nBlocks))+i),int((nBlockssens+nBlocksppi)*(nTapevents/nBlocks)),nBlocksppi)]])/(lattot[number,(nBlockssens+i)]-latnan[number,(nBlockssens+i)])
        towardshalf[number,(nBlockssens+i)] = np.nansum(taptowardshalf[number, [*range(int(((nBlockssens)*(nTapevents/nBlocks))+i),int((nBlockssens+nBlocksppi)*(nTapevents/nBlocks)),nBlocksppi)]])/np.count_nonzero(np.isfinite(taptowardshalf[number, [*range(int(((nBlockssens)*(nTapevents/nBlocks))+i),int((nBlockssens+nBlocksppi)*(nTapevents/nBlocks)),nBlocksppi)]]))

        avgbend[number,(nBlockssens+i)] = np.nanmean(abs(tapbendmax[number, [*range(int(((nBlockssens)*(nTapevents/nBlocks))+i),int((nBlockssens+nBlocksppi)*(nTapevents/nBlocks)),nBlocksppi)]]))
        medbend[number,(nBlockssens+i)] = np.nanmedian(abs(tapbendmax[number, [*range(int(((nBlockssens)*(nTapevents/nBlocks))+i),int((nBlockssens+nBlocksppi)*(nTapevents/nBlocks)),nBlocksppi)]]))
        avgorient[number,(nBlockssens+i)] = np.nanmean(abs(tapdeltaorient[number, [*range(int(((nBlockssens)*(nTapevents/nBlocks))+i),int((nBlockssens+nBlocksppi)*(nTapevents/nBlocks)),nBlocksppi)]]))
        medorient[number,(nBlockssens+i)] = np.nanmedian(abs(tapdeltaorient[number, [*range(int(((nBlockssens)*(nTapevents/nBlocks))+i),int((nBlockssens+nBlocksppi)*(nTapevents/nBlocks)),nBlocksppi)]]))
        avgdisp[number,(nBlockssens+i)] = np.nanmean(abs(tapdisplacement[number, [*range(int(((nBlockssens)*(nTapevents/nBlocks))+i),int((nBlockssens+nBlocksppi)*(nTapevents/nBlocks)),nBlocksppi)]]))
        meddisp[number,(nBlockssens+i)] = np.nanmedian(abs(tapdisplacement[number, [*range(int(((nBlockssens)*(nTapevents/nBlocks))+i),int((nBlockssens+nBlocksppi)*(nTapevents/nBlocks)),nBlocksppi)]]))
        avgdist[number,(nBlockssens+i)] = np.nanmean(abs(tapdistance[number, [*range(int(((nBlockssens)*(nTapevents/nBlocks))+i),int((nBlockssens+nBlocksppi)*(nTapevents/nBlocks)),nBlocksppi)]]))
        meddist[number,(nBlockssens+i)] = np.nanmedian(abs(tapdistance[number, [*range(int(((nBlockssens)*(nTapevents/nBlocks))+i),int((nBlockssens+nBlocksppi)*(nTapevents/nBlocks)),nBlocksppi)]]))
        avgduration[number,(nBlockssens+i)] = np.nanmean(abs(tapduration[number, [*range(int(((nBlockssens)*(nTapevents/nBlocks))+i),int((nBlockssens+nBlocksppi)*(nTapevents/nBlocks)),nBlocksppi)]]))
        medduration[number,(nBlockssens+i)] = np.nanmedian(abs(tapduration[number, [*range(int(((nBlockssens)*(nTapevents/nBlocks))+i),int((nBlockssens+nBlocksppi)*(nTapevents/nBlocks)),nBlocksppi)]]))
        avgangvelmax[number,(nBlockssens+i)] = np.nanmean(abs(angdispmax[number, [*range(int(((nBlockssens)*(nTapevents/nBlocks))+i),int((nBlockssens+nBlocksppi)*(nTapevents/nBlocks)),nBlocksppi)]]))
        medangvelmax[number,(nBlockssens+i)] = np.nanmedian(abs(angdispmax[number, [*range(int(((nBlockssens)*(nTapevents/nBlocks))+i),int((nBlockssens+nBlocksppi)*(nTapevents/nBlocks)),nBlocksppi)]]))
 


#count number of nan and total events and make calculations for hab events
for number in range(nFish):
    for i in range(nBlockshab):
        slcnan[number,(nBlockssens+nBlocksppi+i)] = np.count_nonzero(np.isnan(latencytaps[number, int((nBlockssens+nBlocksppi+i)*(nTapevents/nBlocks)):int(((nBlockssens+nBlocksppi+i)+1)*(nTapevents/nBlocks))])) - np.count_nonzero(np.isfinite(nomvmttaps[number, int((nBlockssens+nBlocksppi+i)*(nTapevents/nBlocks)):int(((nBlockssens+nBlocksppi+i)+1)*(nTapevents/nBlocks))]))
        slctot[number,(nBlockssens+nBlocksppi+i)] = np.size(latencytaps[number, int((nBlockssens+nBlocksppi+i)*(nTapevents/nBlocks)):int(((nBlockssens+nBlocksppi+i)+1)*(nTapevents/nBlocks))])
        latnan[number,(nBlockssens+nBlocksppi+i)] = np.count_nonzero(np.isnan(latencytaps[number, int((nBlockssens+nBlocksppi+i)*(nTapevents/nBlocks)):int(((nBlockssens+nBlocksppi+i)+1)*(nTapevents/nBlocks))]))
        lattot[number,(nBlockssens+nBlocksppi+i)] = np.size(latencytaps[number, int((nBlockssens+nBlocksppi+i)*(nTapevents/nBlocks)):int(((nBlockssens+nBlocksppi+i)+1)*(nTapevents/nBlocks))])
        tapbendmaxnan[number,(nBlockssens+nBlocksppi+i)] = np.count_nonzero(np.isnan(tapbendmax[number, int((nBlockssens+nBlocksppi+i)*(nTapevents/nBlocks)):int(((nBlockssens+nBlocksppi+i)+1)*(nTapevents/nBlocks))]))
        tapbendmaxtot[number,(nBlockssens+nBlocksppi+i)] = np.size(tapbendmax[number, int((nBlockssens+nBlocksppi+i)*(nTapevents/nBlocks)):int(((nBlockssens+nBlocksppi+i)+1)*(nTapevents/nBlocks))])
        tapdeltaorientnan[number,(nBlockssens+nBlocksppi+i)] = np.count_nonzero(np.isnan(tapdeltaorient[number, int((nBlockssens+nBlocksppi+i)*(nTapevents/nBlocks)):int(((nBlockssens+nBlocksppi+i)+1)*(nTapevents/nBlocks))]))
        tapdeltaorienttot[number,(nBlockssens+nBlocksppi+i)] = np.size(tapdeltaorient[number, int((nBlockssens+nBlocksppi+i)*(nTapevents/nBlocks)):int(((nBlockssens+nBlocksppi+i)+1)*(nTapevents/nBlocks))])
        tapdisplacementnan[number,(nBlockssens+nBlocksppi+i)] = np.count_nonzero(np.isnan(tapdisplacement[number, int((nBlockssens+nBlocksppi+i)*(nTapevents/nBlocks)):int(((nBlockssens+nBlocksppi+i)+1)*(nTapevents/nBlocks))]))
        tapdisplacementtot[number,(nBlockssens+nBlocksppi+i)] = np.size(tapdisplacement[number, int((nBlockssens+nBlocksppi+i)*(nTapevents/nBlocks)):int(((nBlockssens+nBlocksppi+i)+1)*(nTapevents/nBlocks))])
        tapdistancenan[number,(nBlockssens+nBlocksppi+i)] = np.count_nonzero(np.isnan(tapdistance[number, int((nBlockssens+nBlocksppi+i)*(nTapevents/nBlocks)):int(((nBlockssens+nBlocksppi+i)+1)*(nTapevents/nBlocks))]))
        tapdistancetot[number,(nBlockssens+nBlocksppi+i)] = np.size(tapdistance[number, int((nBlockssens+nBlocksppi+i)*(nTapevents/nBlocks)):int(((nBlockssens+nBlocksppi+i)+1)*(nTapevents/nBlocks))])
        angdispmaxnan[number,(nBlockssens+nBlocksppi+i)] = np.count_nonzero(np.isnan(angdispmax[number, int((nBlockssens+nBlocksppi+i)*(nTapevents/nBlocks)):int(((nBlockssens+nBlocksppi+i)+1)*(nTapevents/nBlocks))]))
        angdispmaxtot[number,(nBlockssens+nBlocksppi+i)] = np.size(angdispmax[number, int((nBlockssens+nBlocksppi+i)*(nTapevents/nBlocks)):int(((nBlockssens+nBlocksppi+i)+1)*(nTapevents/nBlocks))])


        right[number,(nBlockssens+nBlocksppi+i)] = np.nansum(tapright[number, int((nBlockssens+nBlocksppi+i)*(nTapevents/nBlocks)):int(((nBlockssens+nBlocksppi+i)+1)*(nTapevents/nBlocks))])/(lattot[number,(nBlockssens+nBlocksppi+i)]-latnan[number,(nBlockssens+nBlocksppi+i)])
        towards[number,(nBlockssens+nBlocksppi+i)] = np.nansum(taptowards[number, int((nBlockssens+nBlocksppi+i)*(nTapevents/nBlocks)):int(((nBlockssens+nBlocksppi+i)+1)*(nTapevents/nBlocks))])/(lattot[number,(nBlockssens+nBlocksppi+i)]-latnan[number,(nBlockssens+nBlocksppi+i)])
        towardshalf[number,(nBlockssens+nBlocksppi+i)] = np.nansum(taptowardshalf[number, int((nBlockssens+nBlocksppi+i)*(nTapevents/nBlocks)):int(((nBlockssens+nBlocksppi+i)+1)*(nTapevents/nBlocks))])/np.count_nonzero(np.isfinite(taptowardshalf[number, int((nBlockssens+nBlocksppi+i)*(nTapevents/nBlocks)):int(((nBlockssens+nBlocksppi+i)+1)*(nTapevents/nBlocks))]))

        
        avglat[number,(nBlockssens+nBlocksppi+i)] = np.nanmean(latencytapsms[number, int((nBlockssens+nBlocksppi+i)*(nTapevents/nBlocks)):int(((nBlockssens+nBlocksppi+i)+1)*(nTapevents/nBlocks))])
        medlat[number,(nBlockssens+nBlocksppi+i)] = np.nanmedian(latencytapsms[number, int((nBlockssens+nBlocksppi+i)*(nTapevents/nBlocks)):int(((nBlockssens+nBlocksppi+i)+1)*(nTapevents/nBlocks))])
        slc[number,(nBlockssens+nBlocksppi+i)] = np.nansum(slctaps[number, int((nBlockssens+nBlocksppi+i)*(nTapevents/nBlocks)):int(((nBlockssens+nBlocksppi+i)+1)*(nTapevents/nBlocks))])/(slctot[number,(nBlockssens+nBlocksppi+i)]-slcnan[number,(nBlockssens+nBlocksppi+i)])
        llc[number,(nBlockssens+nBlocksppi+i)] = np.nansum(llctaps[number, int((nBlockssens+nBlocksppi+i)*(nTapevents/nBlocks)):int(((nBlockssens+nBlocksppi+i)+1)*(nTapevents/nBlocks))])/(slctot[number,(nBlockssens+nBlocksppi+i)]-slcnan[number,(nBlockssens+nBlocksppi+i)])
        react[number,(nBlockssens+nBlocksppi+i)] = np.nansum(reacttaps[number, int((nBlockssens+nBlocksppi+i)*(nTapevents/nBlocks)):int(((nBlockssens+nBlocksppi+i)+1)*(nTapevents/nBlocks))])/(slctot[number,(nBlockssens+nBlocksppi+i)]-slcnan[number,(nBlockssens+nBlocksppi+i)])
        nomvmt[number,(nBlockssens+nBlocksppi+i)] = np.nansum(nomvmttaps[number, int((nBlockssens+nBlocksppi+i)*(nTapevents/nBlocks)):int(((nBlockssens+nBlocksppi+i)+1)*(nTapevents/nBlocks))])/(slctot[number,(nBlockssens+nBlocksppi+i)]-slcnan[number,(nBlockssens+nBlocksppi+i)])
        avgbend[number,(nBlockssens+nBlocksppi+i)] = np.nanmean(abs(tapbendmax[number, int((nBlockssens+nBlocksppi+i)*(nTapevents/nBlocks)):int(((nBlockssens+nBlocksppi+i)+1)*(nTapevents/nBlocks))]))
        medbend[number,(nBlockssens+nBlocksppi+i)] = np.nanmedian(abs(tapbendmax[number, int((nBlockssens+nBlocksppi+i)*(nTapevents/nBlocks)):int(((nBlockssens+nBlocksppi+i)+1)*(nTapevents/nBlocks))]))
        avgorient[number,(nBlockssens+nBlocksppi+i)] = np.nanmean(abs(tapdeltaorient[number, int((nBlockssens+nBlocksppi+i)*(nTapevents/nBlocks)):int(((nBlockssens+nBlocksppi+i)+1)*(nTapevents/nBlocks))]))
        medorient[number,(nBlockssens+nBlocksppi+i)] = np.nanmedian(abs(tapdeltaorient[number, int((nBlockssens+nBlocksppi+i)*(nTapevents/nBlocks)):int(((nBlockssens+nBlocksppi+i)+1)*(nTapevents/nBlocks))]))
        avgdisp[number,(nBlockssens+nBlocksppi+i)] = np.nanmean(abs(tapdisplacement[number, int((nBlockssens+nBlocksppi+i)*(nTapevents/nBlocks)):int(((nBlockssens+nBlocksppi+i)+1)*(nTapevents/nBlocks))]))
        meddisp[number,(nBlockssens+nBlocksppi+i)] = np.nanmedian(abs(tapdisplacement[number, int((nBlockssens+nBlocksppi+i)*(nTapevents/nBlocks)):int(((nBlockssens+nBlocksppi+i)+1)*(nTapevents/nBlocks))]))
        avgdist[number,(nBlockssens+nBlocksppi+i)] = np.nanmean(abs(tapdistance[number, int((nBlockssens+nBlocksppi+i)*(nTapevents/nBlocks)):int(((nBlockssens+nBlocksppi+i)+1)*(nTapevents/nBlocks))]))
        meddist[number,(nBlockssens+nBlocksppi+i)] = np.nanmedian(abs(tapdistance[number, int((nBlockssens+nBlocksppi+i)*(nTapevents/nBlocks)):int(((nBlockssens+nBlocksppi+i)+1)*(nTapevents/nBlocks))]))
        avgduration[number,(nBlockssens+nBlocksppi+i)] = np.nanmean(abs(tapduration[number, int((nBlockssens+nBlocksppi+i)*(nTapevents/nBlocks)):int(((nBlockssens+nBlocksppi+i)+1)*(nTapevents/nBlocks))]))
        medduration[number,(nBlockssens+nBlocksppi+i)] = np.nanmedian(abs(tapduration[number, int((nBlockssens+nBlocksppi+i)*(nTapevents/nBlocks)):int(((nBlockssens+nBlocksppi+i)+1)*(nTapevents/nBlocks))]))
        avgangvelmax[number,(nBlockssens+nBlocksppi+i)] = np.nanmean(abs(angdispmax[number, int((nBlockssens+nBlocksppi+i)*(nTapevents/nBlocks)):int(((nBlockssens+nBlocksppi+i)+1)*(nTapevents/nBlocks))]))
        medangvelmax[number,(nBlockssens+nBlocksppi+i)] = np.nanmedian(abs(angdispmax[number, int((nBlockssens+nBlocksppi+i)*(nTapevents/nBlocks)):int(((nBlockssens+nBlocksppi+i)+1)*(nTapevents/nBlocks))]))


  
print("Saving tracking data arrays....")
np.savez(inDir + 'Results/TrackedVideos/' + Name + '/AnalyzedData.npz', medfiltcentY=medfiltcentY, medfiltcentX=medfiltcentX, disp=disp, filtdisp=filtdisp, 
         latencytaps=latencytaps, latencytapsms=latencytapsms, slctaps=slctaps, 
         llctaps=llctaps, reacttaps=reacttaps, nomvmttaps=nomvmttaps, tapbendmax=tapbendmax,
         tapdeltaorient=tapdeltaorient, tapdisplacement=tapdisplacement, tapdistance=tapdistance, movementyesno=movementyesno,
         movementyesnofill=movementyesnofill,
         slcnan=slcnan, slctot=slctot, latnan=latnan, lattot=lattot, tapbendmaxnan=tapbendmaxnan, tapbendmaxtot=tapbendmaxtot, 
         angdispmaxnan=angdispmaxnan, angdispmaxtot=angdispmaxtot, avgangvelmax=avgangvelmax, medangvelmax=medangvelmax,
         tapdeltaorientnan=tapdeltaorientnan, tapdeltaorienttot=tapdeltaorienttot, tapdisplacementnan=tapdisplacementnan, tapdisplacementtot=tapdisplacementtot, 
         tapdistancenan=tapdistancenan, tapdistancetot=tapdistancetot, right=right, towards=towards, towardshalf=towardshalf,
         avglat=avglat, medlat=medlat, slc=slc, llc=llc, react=react, nomvmt=nomvmt, avgbend=avgbend, medbend=medbend,
         avgorient=avgorient, medorient=medorient, avgdisp=avgdisp, meddisp=meddisp, avgdist=avgdist, meddist=meddist, avgduration=avgduration, medduration=medduration
         )
```
