## Supplemental Zip file 1 - python tracking scripts for "Mitochondrial genes in the 22q11.2 deleted region regulate neural stem and progenitor cell proliferation": VMR tracking file.py

```
# -*- coding: utf-8 -*-
"""
VMR Tracking code

by Philip D. Campbell


"""

import numpy as np
import cv2
#from TrackedObject import TrackedObject as TO
import matplotlib.pyplot as plt
import os
import re #regular expressions library
from skimage.measure import label, regionprops, find_contours
from skimage.morphology import remove_small_objects
from skimage import img_as_ubyte
import scipy.misc
import scipy.interpolate
import scipy.io
import pims
from tqdm import tqdm
import pandas as pd
import math
import imageio


import scipy.stats as stats
import seaborn as sns
from matplotlib.pyplot import figure
from sklearn import metrics 


#set directory containing image files
inDir = ""

# names of videos to track
Name = ''

#set gene name
Gene = ""

# set directory containing background files; background video used for VMR assay is itself
inDirbg = ""

# names of videos for background
Namebg = ''


#set if Tracking images to be saved
SaveTrackingIm = True; # or False

#set locations
desttracked = inDir + 'Results/TrackedVideos/' + Name + '/trackedimages/'
desttracking = inDir + 'Results/TrackedVideos/' + Name + '/'
destgene = inDir + 'Results/' + Gene + '/' + Name + '/'

# Number of fish to track
nFish = 100

#Number of rows and columns of wells
nRow = 10
nCol = 10

#Number of pixels in x and y direction
nPixel = 1024

#frame rate in Hz
framerate = 20

# time between frames in milliseconds
framelength = 1/framerate * 1000

# Define block time in seconds; ie bins you would like data to be analyzed in
blocksec = 60

# Define number of total blocks
nBlocks = 16

#if no folder, make one
os.makedirs(os.path.dirname(desttracked), exist_ok=True)
os.makedirs(os.path.dirname(desttracking), exist_ok=True)
os.makedirs(os.path.dirname(destgene), exist_ok=True)

#load video
video = pims.PyAVReaderIndexed(inDir + Name)

# number of frames of video
nFrames = len(video)

# number of frames per block
nFramesblock = math.floor((nFrames-5)/nBlocks)


#load background video
videobg = pims.PyAVReaderIndexed(inDirbg + Namebg)

# number of frames of video
nFramesbg = len(videobg)

#background subtraction
#first 5 frames should be used to create background - These are acquired in 10mins of aclimation, spread out over the 10mins
fr1 = videobg[0][:,:,0]
fr2 = videobg[1][:,:,0]
fr3 = videobg[2][:,:,0]
fr4 = videobg[3][:,:,0]
fr5 = videobg[4][:,:,0]
fr6 = videobg[5][:,:,0]

bg1 = np.median(np.stack((fr1, fr2, fr3, fr4, fr5, fr6), axis=2), axis=2)

bg = cv2.GaussianBlur(bg1.astype(np.float64), (3,3),0)


def tracking(bg, img, index):
 
    #Do a background subtraction, and blur the background
    mask = abs(bg - img)

    mask_smooth = cv2.GaussianBlur(mask, (3,3),0)

    #min_val = -50.0
    #img_crop = np.where(mask < min_val, img_crop, np.zeros_like(img_crop))
    
    #threshold the mask
    th_img = cv2.threshold(mask_smooth, 15, 255, cv2.THRESH_BINARY)[1]
    
    #label the connected components in the thresholded mask
    lb = label(th_img)
    
    #remove small objects from thresholded image
    lb_sorem = remove_small_objects(lb, 40)
    
    #re-label since small objects removes so get 1 to nFish
    lb2 = label(lb_sorem)

    #get properties of objects
    props = regionprops(lb2)

    # of objects
    totobj = np.amax(lb2)
    print(str(totobj) + "in" + str(index+1))

    
    #copy the smoothed mask into a temporary image
    im_tmp = np.copy(mask_smooth)
    

    #empty arrays to hold centroid values    
    y_cent = np.zeros(nFish)
    y_cent[:] = np.nan
    x_cent = np.zeros(nFish)
    x_cent[:] = np.nan


    #Loop through all objects    
    for ind in range(0, totobj):       
        
        #find the centroid of fish      
        cent = props[ind].centroid
        
        #reset fishrow and fishcol tracker
        fishrow=0
        fishcol=0
        
        #go through and get row and column of this fish so can use it to index array; fish numbered from 1 in top left to nFish in bottom right
        for row in range(1, nRow+1):

            if cent[0] < row*nPixel/nRow:
                fishrow = row
                break
        
        for col in range(1, nCol+1):
            if cent[1] < col*nPixel/nCol:
                fishcol = col
                break
        
        #this will return number of fish from 1 to nFish starting in top left corner with 1 and going L to R
        fishnum = ((fishrow-1) * nCol) + fishcol
        
        #if there is already data in arrays for this fish number, put NaN for values and go to top of for loop for next object     
        if np.isfinite(y_cent[fishnum-1]):
            y_cent[fishnum-1] = np.nan
            x_cent[fishnum-1] = np.nan
            continue
        
        #find the centroid of the largest component
        y_cent[fishnum-1], x_cent[fishnum-1] = cent

      
        #set the cent point to 255 in temp image
        im_tmp[y_cent[fishnum-1].astype(int), x_cent[fishnum-1].astype(int)] = 255;


    return im_tmp, y_cent, x_cent


#Initialize arrays to hold values
cents = np.zeros((nFish, 2, nFrames))

#tracking

print("Tracking....")
    
#tqdm shows a progress bar around any iterable
for index, img in enumerate(tqdm(video)):
       
        if (index < nFrames) and (index >= 5):

            ind = index-5
            
            img = img[:,:,0] # its actually a black and white image, but gets read in as three channel
            
            # blur image
            img = cv2.GaussianBlur(img.astype(np.float64), (3,3),0)
            
            #calling the tracking image with bg and individual img
            timg, y_cent, x_cent = tracking(bg, img, ind)
            
            cents[:, 0, ind] = y_cent
            cents[:, 1, ind] = x_cent

            #save the tracked coordinates and the tracked image files
            if SaveTrackingIm:
                
                cv2.imwrite(inDir + 'Results/TrackedVideos/' + Name + '/trackedimages/' + Name + '_' + str(index) +'.png', np.vstack((timg)))        
        

print("Saving tracking data arrays....")
np.savez(inDir + 'Results/TrackedVideos/' + Name + '/TrackData.npz', cents=cents)

        
import seaborn as sns
from scipy.signal import medfilt

# START CALCULATIONS HERE

#setup med-filtered arrays
medfiltcentY = np.zeros((nFish, nFrames))
medfiltcentY[:,:] = np.nan
medfiltcentX = np.zeros((nFish, nFrames))
medfiltcentX[:,:] = np.nan


for number in range(nFish):
        medfiltcentY[number,:] = medfilt(cents[number,0,:],3)
        medfiltcentX[number,:] = medfilt(cents[number,1,:],3)
      
#set up arrays for data
disp = np.zeros((nFish, nFrames-1))
disp[:,:] = np.nan
filtdisp = np.zeros((nFish, nFrames-1))
filtdisp[:,:] = np.nan


thigmo = np.zeros((nFish, nFrames))
thigmo[:,:] = np.nan
filtthigmo = np.zeros((nFish, nFrames))
filtthigmo[:,:] = np.nan

centeryesno = np.zeros((nFish, nFrames))
centeryesno[:,:] = np.nan

movementyesno = np.zeros((nFish, nFrames-1))
movementyesno[:,:] = np.nan
movementyesnofill = np.zeros((nFish, nFrames-1))
movementyesnofill[:,:] = np.nan


boutsperblock= np.zeros((nFish, nBlocks))
boutsperblock[:,:] = np.nan
totdistperblock = np.zeros((nFish, nBlocks))
totdistperblock[:,:] = np.nan
totdispperblock = np.zeros((nFish, nBlocks))
totdispperblock[:,:] = np.nan
tottimemovingperblock = np.zeros((nFish, nBlocks))
tottimemovingperblock[:,:] = np.nan
totavgspeedperblock = np.zeros((nFish, nBlocks))
totavgspeedperblock[:,:] = np.nan
avgthigmoperblock = np.zeros((nFish, nBlocks))
avgthigmoperblock[:,:] = np.nan
medthigmoperblock = np.zeros((nFish, nBlocks))
medthigmoperblock[:,:] = np.nan
fractionouterperblock = np.zeros((nFish, nBlocks))
fractionouterperblock[:,:] = np.nan

boutsperperiod= np.zeros((nFish, 2))
boutsperperiod[:,:] = np.nan
totdistperperiod = np.zeros((nFish, 2))
totdistperperiod[:,:] = np.nan
totdispperperiod = np.zeros((nFish, 2))
totdispperperiod[:,:] = np.nan
tottimemovingperperiod = np.zeros((nFish, 2))
tottimemovingperperiod[:,:] = np.nan
totavgspeedperperiod = np.zeros((nFish, 2))
totavgspeedperperiod[:,:] = np.nan
avgthigmoperperiod = np.zeros((nFish, 2))
avgthigmoperperiod[:,:] = np.nan
medthigmoperperiod = np.zeros((nFish, 2))
medthigmoperperiod[:,:] = np.nan
fractionouterperperiod = np.zeros((nFish, 2))
fractionouterperperiod[:,:] = np.nan

avgdistperboutperblock = np.zeros((nFish, nBlocks))
avgdistperboutperblock[:,:] = np.nan
avgdispperboutperblock = np.zeros((nFish, nBlocks))
avgdispperboutperblock[:,:] = np.nan
avgtimemovingperboutperblock = np.zeros((nFish, nBlocks))
avgtimemovingperboutperblock[:,:] = np.nan
avgspeedperboutperblock = np.zeros((nFish, nBlocks))
avgspeedperboutperblock[:,:] = np.nan

meddistperboutperblock = np.zeros((nFish, nBlocks))
meddistperboutperblock[:,:] = np.nan
meddispperboutperblock = np.zeros((nFish, nBlocks))
meddispperboutperblock[:,:] = np.nan
medtimemovingperboutperblock = np.zeros((nFish, nBlocks))
medtimemovingperboutperblock[:,:] = np.nan
medspeedperboutperblock = np.zeros((nFish, nBlocks))
medspeedperboutperblock[:,:] = np.nan

avgdistperboutperperiod = np.zeros((nFish, 2))
avgdistperboutperperiod[:,:] = np.nan
avgdispperboutperperiod = np.zeros((nFish, 2))
avgdispperboutperperiod[:,:] = np.nan
avgtimemovingperboutperperiod = np.zeros((nFish, 2))
avgtimemovingperboutperperiod[:,:] = np.nan
avgspeedperboutperperiod = np.zeros((nFish, 2))
avgspeedperboutperperiod[:,:] = np.nan

meddistperboutperperiod = np.zeros((nFish, 2))
meddistperboutperperiod[:,:] = np.nan
meddispperboutperperiod = np.zeros((nFish, 2))
meddispperboutperperiod[:,:] = np.nan
medtimemovingperboutperperiod = np.zeros((nFish, 2))
medtimemovingperboutperperiod[:,:] = np.nan
medspeedperboutperperiod = np.zeros((nFish, 2))
medspeedperboutperperiod[:,:] = np.nan

distbouts = np.zeros((nFish, nBlocks,1000))
distbouts[:,:,:] = np.nan
dispbouts = np.zeros((nFish, nBlocks,1000))
dispbouts[:,:,:] = np.nan
timebouts = np.zeros((nFish, nBlocks,1000))
timebouts[:,:,:] = np.nan
speedbouts = np.zeros((nFish, nBlocks,1000))
speedbouts[:,:,:] = np.nan


print("Calculating displacement and thigmotaxis....")
# calculating displacement and thigmotaxis
for number in range(nFish):
    i = 0
    while i < nFrames:
        block = math.floor(i/nFramesblock)
        if  np.isnan(cents[number, 0, (block*nFramesblock):((block+1)*nFramesblock)]).any():
            i=(block+1)*nFramesblock
            #if no centroid is defined in any frame in this block, skip to next block
            continue

        else:            
            #calculate thigmotaxis for each frame
            row = math.ceil((number+1)/nCol)
            col = (number+1) - (nCol*(row-1))
            ycenter = (nPixel/nRow)/2 + ((row - 1)*(nPixel/nRow))
            xcenter = (nPixel/nCol)/2 + ((col - 1)*(nPixel/nCol))
            thigmo[number,i] = np.sqrt((cents[number, 0, i]-ycenter)**2 + (cents[number, 1, i]-xcenter)**2)
            filtthigmo[number,i] = np.sqrt((medfiltcentY[number, i]-ycenter)**2 + (medfiltcentX[number, i]-xcenter)**2)

            if thigmo[number,i]>30:
                centeryesno[number,i] = 1
            if thigmo[number,i]<=30:
                centeryesno[number,i] = 0

            #calculate displacement between each frame
            if i>0:
                disp[number,i-1] = np.sqrt((cents[number, 0, i]-cents[number, 0, i-1])**2 + (cents[number, 1, i]-cents[number, 1, i-1])**2) 
                filtdisp[number,i-1] = np.sqrt((medfiltcentY[number, i]-medfiltcentY[number, i-1])**2 + (medfiltcentX[number, i]-medfiltcentX[number, i-1])**2) 
                   # define movement as displacement >0.2 and populate array with 1 for >0.2 and 0 for <0.2; nan if displ nan
                if disp[number,i-1] > 0.7:
                    movementyesno[number,i-1] = 1
                else:
                    movementyesno[number,i-1] = 0
            
            i=i+1


print("Defining movement....")
#Attempt to fill holes of movement array. Will eliminate singleton 1s or 0s so that for movement will need >1 frame with value of
                       #1 and to stop movement will need >1 frame with value of 0
# fill initial frame with values of initial array
movementyesnofill[:, 0] = movementyesno[:, 0]

for number in range(nFish):
    for i in range(1,nFrames-1):
        if  np.isnan(movementyesno[number, i]):
            continue
        #if nan either before or after value i, just keep value because cannot tell
        if  np.isnan(movementyesnofill[number, i-1]):
            movementyesnofill[number, i] = movementyesno[number, i]
            continue
        if i < nFrames-2:
            if np.isnan(movementyesno[number, i+1]):
                movementyesnofill[number, i] = movementyesno[number, i]
                continue


# here is where we define movement and stopping as >1 frame of either 1 or 0
        if movementyesno[number, i] == 0:
            if movementyesnofill[number, i-1] == 0:
                movementyesnofill[number, i] = 0
            if movementyesnofill[number, i-1] == 1:
                if i < nFrames-2:
                    if movementyesno[number, i+1] == 1:
                        movementyesnofill[number, i] = 1
                    if movementyesno[number, i+1] == 0:
                        movementyesnofill[number, i] = 0
                else:
                    movementyesnofill[number, i] = 0
        
        if movementyesno[number, i] == 1:
            if movementyesnofill[number, i-1] == 1:
                movementyesnofill[number, i] = 1
            if movementyesnofill[number, i-1] == 0:
                if i < nFrames-2:
                    if movementyesno[number, i+1] == 0:
                        movementyesnofill[number, i] = 0
                    if movementyesno[number, i+1] == 1:
                        movementyesnofill[number, i] = 1
                else:
                    movementyesnofill[number, i] = 1


print("Calculating values per block and per period")
for number in range(nFish):
    i=0
    while i < nBlocks:
        #check if no cent values anywhere, if so, skip this block
        if  np.isnan(cents[number, 0, (i*nFramesblock):((i+1)*nFramesblock)]).any():
            i=i+1
            #if no centroid is defined in any frame in this block, skip to next block
            continue        

        else:
            #label bouts in block
            lb = label(movementyesnofill[number, (i*nFramesblock):(((i+1)*nFramesblock)-1)])
            
            #remove bouts less than 2 - though shouldn't be any anyways
            lb_sorem = remove_small_objects(lb, 2)
            
            #re-label since small objects removed
            lb2 = label(lb_sorem)
            
            # Number of bouts for this block
            boutsperblock[number,i] = np.amax(lb2)
            
            #define array of distances in this block
            x = disp[number, (i*nFramesblock):(((i+1)*nFramesblock)-1)]
            
            #total distance traveled during periods defined as movements
            totdistperblock[number, i] = np.sum(x[lb2!=0])
   
            #total displacement for this block
            totdispperblock[number,i] = np.sqrt((cents[number, 0, (i*nFramesblock)]-cents[number, 0, (((i+1)*nFramesblock)-1)])**2 + (cents[number, 1, (i*nFramesblock)]-cents[number, 1, (((i+1)*nFramesblock)-1)])**2)
          
            #time moving for this block in ms
            tottimemovingperblock[number,i] = np.sum(movementyesnofill[number,(i*nFramesblock):(((i+1)*nFramesblock)-1)])*framelength
            
            #average speed for entire block pix/s
            totavgspeedperblock[number,i] = (totdistperblock[number, i]/tottimemovingperblock[number,i])*1000
         
            #average and median distance from center for entire block
            avgthigmoperblock[number,i] = np.nanmean(thigmo[number, (i*nFramesblock):((i+1)*nFramesblock)])
            medthigmoperblock[number,i] = np.nanmedian(thigmo[number, (i*nFramesblock):((i+1)*nFramesblock)])
            
            # fraction of time in outer part of well vs center
            fractionouterperblock[number,i] = np.sum(centeryesno[number, (i*nFramesblock):((i+1)*nFramesblock)])/nFramesblock
           
            #calulating values per bout
            
            index = 0

            while index < int(boutsperblock[number,i]):
                #distance travelled of this bout
                distbouts[number,i,index] = np.sum(x[lb2==(index+1)])
                
                #define first and last frame of this bout
                first = int(np.where(lb2==(index+1))[0][0])
                last = int(np.where(lb2==(index+1))[0][-1]+1)
                
                # displacement of this bout
                dispbouts[number,i,index] = np.sqrt((cents[number, 0, ((i*nFramesblock)+first)]-cents[number, 0, ((i*nFramesblock)+last)])**2 + (cents[number, 1, ((i*nFramesblock)+first)]-cents[number, 1, ((i*nFramesblock)+last)])**2)
                
                # time moving of this bout
                timebouts[number,i,index] = np.count_nonzero(lb2 == (index+1))*framelength
                
                #average speed of this bout pix/s
                speedbouts[number,i,index] = (distbouts[number,i,index]/timebouts[number,i,index])*1000
                
                index = index+1         
            
            
            #calculating perbout, per block values
                        
            avgdistperboutperblock[number, i] = np.nanmean(distbouts[number,i,:])
            meddistperboutperblock[number, i] = np.nanmedian(distbouts[number,i,:])
            
            avgdispperboutperblock[number, i] = np.nanmean(dispbouts[number,i,:])
            meddispperboutperblock[number, i] = np.nanmedian(dispbouts[number,i,:])
            
            avgtimemovingperboutperblock[number, i] = np.nanmean(timebouts[number,i,:])
            medtimemovingperboutperblock[number, i] = np.nanmedian(timebouts[number,i,:])
            
            avgspeedperboutperblock[number, i] = np.nanmean(speedbouts[number,i,:])
            medspeedperboutperblock[number, i] = np.nanmedian(speedbouts[number,i,:])
                     
            i=i+1

#calulating total for entire period (light is index 0, dark is index 1)    
    boutsperperiod[number,0] = np.sum(boutsperblock[number,:int(nBlocks/2)])
    boutsperperiod[number,1] = np.sum(boutsperblock[number,int(nBlocks/2):])
    
    totdistperperiod[number,0] = np.sum(totdistperblock[number,:int(nBlocks/2)])
    totdistperperiod[number,1] = np.sum(totdistperblock[number,int(nBlocks/2):])

    totdispperperiod[number,0] = np.sum(totdispperblock[number,:int(nBlocks/2)])
    totdispperperiod[number,1] = np.sum(totdispperblock[number,int(nBlocks/2):])

    tottimemovingperperiod[number,0] = np.sum(tottimemovingperblock[number,:int(nBlocks/2)])
    tottimemovingperperiod[number,1] = np.sum(tottimemovingperblock[number,int(nBlocks/2):])

    totavgspeedperperiod[number,0] = (totdistperperiod[number,0]/tottimemovingperperiod[number,0])*1000
    totavgspeedperperiod[number,1] = (totdistperperiod[number,1]/tottimemovingperperiod[number,1])*1000


    avgthigmoperperiod[number,0] = np.nanmean(thigmo[number, :(int(nBlocks/2)*nFramesblock)])
    avgthigmoperperiod[number,1] = np.nanmean(thigmo[number, (int(nBlocks/2)*nFramesblock):])                                                    

    medthigmoperperiod[number,0] = np.nanmedian(thigmo[number, :(int(nBlocks/2)*nFramesblock)])
    medthigmoperperiod[number,1] = np.nanmedian(thigmo[number, (int(nBlocks/2)*nFramesblock):])  
    
    fractionouterperperiod[number,0] = np.sum(centeryesno[number, :(int(nBlocks/2)*nFramesblock)])/(nFramesblock*(int(nBlocks/2)))
    fractionouterperperiod[number,1] = np.sum(centeryesno[number, (int(nBlocks/2)*nFramesblock):])/(nFramesblock*(int(nBlocks/2)))
        

#calculating per bout, per period values
    avgdistperboutperperiod[number,0] = np.nanmean(distbouts[number,:int(nBlocks/2),:])
    avgdistperboutperperiod[number,1] = np.nanmean(distbouts[number,int(nBlocks/2):,:])

    meddistperboutperperiod[number,0] = np.nanmedian(distbouts[number,:int(nBlocks/2),:])
    meddistperboutperperiod[number,1] = np.nanmedian(distbouts[number,int(nBlocks/2):,:])

    avgdispperboutperperiod[number,0] = np.nanmean(dispbouts[number,:int(nBlocks/2),:])
    avgdispperboutperperiod[number,1] = np.nanmean(dispbouts[number,int(nBlocks/2):,:])

    meddispperboutperperiod[number,0] = np.nanmedian(dispbouts[number,:int(nBlocks/2),:])
    meddispperboutperperiod[number,1] = np.nanmedian(dispbouts[number,int(nBlocks/2):,:])

    avgtimemovingperboutperperiod[number,0] = np.nanmean(timebouts[number,:int(nBlocks/2),:])
    avgtimemovingperboutperperiod[number,1] = np.nanmean(timebouts[number,int(nBlocks/2):,:])

    medtimemovingperboutperperiod[number,0] = np.nanmedian(timebouts[number,:int(nBlocks/2),:])
    medtimemovingperboutperperiod[number,1] = np.nanmedian(timebouts[number,int(nBlocks/2):,:])

    avgspeedperboutperperiod[number,0] = np.nanmean(speedbouts[number,:int(nBlocks/2),:])
    avgspeedperboutperperiod[number,1] = np.nanmean(speedbouts[number,int(nBlocks/2):,:])

    medspeedperboutperperiod[number,0] = np.nanmedian(speedbouts[number,:int(nBlocks/2),:])
    medspeedperboutperperiod[number,1] = np.nanmedian(speedbouts[number,int(nBlocks/2):,:])

    
  
print("Saving tracking data arrays....")
np.savez(inDir + 'Results/TrackedVideos/' + Name + '/AnalyzedData.npz', medfiltcentY=medfiltcentY, medfiltcentX=medfiltcentX, disp=disp, filtdisp=filtdisp, thigmo=thigmo, filtthigmo=filtthigmo, 
         centeryesno=centeryesno, movementyesno=movementyesno, movementyesnofill=movementyesnofill,
         boutsperblock=boutsperblock, totdistperblock=totdistperblock, totdispperblock=totdispperblock, tottimemovingperblock=tottimemovingperblock,
         totavgspeedperblock=totavgspeedperblock, avgthigmoperblock=avgthigmoperblock, medthigmoperblock=medthigmoperblock, fractionouterperblock=fractionouterperblock,
         boutsperperiod=boutsperperiod, totdistperperiod=totdistperperiod, totdispperperiod=totdispperperiod, tottimemovingperperiod=tottimemovingperperiod, 
         totavgspeedperperiod=totavgspeedperperiod, avgthigmoperperiod=avgthigmoperperiod, medthigmoperperiod=medthigmoperperiod, fractionouterperperiod=fractionouterperperiod,
         avgdistperboutperblock=avgdistperboutperblock, avgdispperboutperblock=avgdispperboutperblock, avgtimemovingperboutperblock=avgtimemovingperboutperblock,
         avgspeedperboutperblock=avgspeedperboutperblock, meddistperboutperblock=meddistperboutperblock, meddispperboutperblock=meddispperboutperblock,
         medtimemovingperboutperblock=medtimemovingperboutperblock, medspeedperboutperblock=medspeedperboutperblock, avgdistperboutperperiod=avgdistperboutperperiod,
         avgdispperboutperperiod=avgdispperboutperperiod, avgtimemovingperboutperperiod=avgtimemovingperboutperperiod, avgspeedperboutperperiod=avgspeedperboutperperiod,
         meddistperboutperperiod=meddistperboutperperiod, meddispperboutperperiod=meddispperboutperperiod, medtimemovingperboutperperiod=medtimemovingperboutperperiod,
         medspeedperboutperperiod=medspeedperboutperperiod, distbouts=distbouts, dispbouts=dispbouts, timebouts=timebouts, speedbouts=speedbouts
         )
```
